# Dynamic interactions between the RNA chaperone Hfq, small regulatory RNAs and mRNAs in live bacterial cells

**DOI:** 10.1101/2020.01.13.903641

**Authors:** Seongjin Park, Karine Prévost, Emily M. Heideman, Marie-Claude Carrier, Matthew A. Reyer, Wei Liu, Eric Massé, Jingyi Fei

**Author notes:** Equal contribution (Seongjin Park, Karine Prévost). Correspondence (JF, Lead contact), (EM).

## Abstract

RNA binding proteins play myriad roles in controlling and regulating RNAs and RNA-mediated functions, often through simultaneous binding to other cellular factors. In bacteria, the RNA chaperone Hfq modulates post-transcriptional gene regulation. Absence of Hfq leads to the loss of fitness and compromises the virulence of bacterial pathogens. Using live-cell super-resolution imaging, we are able to distinguish Hfq binding to different sizes of cellular RNAs. We demonstrate that under normal growth conditions, Hfq exhibits widespread mRNA binding activity. Particularly, the distal face of Hfq contributes mostly to the mRNA binding *in vivo*. In addition, binding of Hfq to these mRNAs can recruit RNase E to promote turnover of these mRNAs in an sRNA-independent manner, providing one mechanism to release Hfq from the pre-bound mRNAs. Finally, our data indicate that sRNAs, once expressed, can either co-occupy Hfq with the mRNA or displace the mRNA from Hfq, suggesting mechanisms through which sRNAs rapidly access Hfq to induce sRNA-mediated gene regulation. Our data collectively demonstrate that Hfq dynamically changes its interactions with different RNAs in response to changes in cellular conditions.

## Main Text

In all three kingdoms of life, RNA binding proteins (RBPs) play myriad roles in controlling and regulating RNAs and RNA-mediated functions. As one of the most abundant RNA binding proteins in bacterial cells, Hfq is an important and prevalent post-transcriptional gene regulator^1–3^. The most recognized functions of Hfq as a chaperone of small regulatory RNA (sRNA)-mediated gene regulation are to stabilize sRNAs and to promote sRNAs binding to their cognate target mRNAs^1–3^. Binding of sRNAs to target mRNAs further leads to changes in the translation activity and the stability of the mRNAs^4, 5^. Moreover, other functions of Hfq in regulating translation and degradation of mRNAs independent of sRNA-mediated regulatory pathway have also been reported^6–10^. The functional importance of Hfq is evident by the fact that loss of Hfq compromises the fitness of bacterial cells, especially under harsh conditions, and abolishes the virulence of bacterial pathogens^11, 12^.

Hfq binds broadly to cellular mRNAs and sRNAs^13–15^, in line with its main biological functions. Hfq can bind RNAs through multiple interfaces of its homohexameric structure. The surface containing the N-terminal α-helices is referred as the “proximal face” of the Hfq hexamer, whereas the opposite surface is referred as the “distal face”, and the outer ring as the “rim” (Figure 1a). The proximal face binds preferably U-rich sequences, and the distal face prefers A-rich sequences, with the exact motif of the A-rich sequence depending on the species^16–20^. The rim can also interact with UA-rich RNAs through the patch of positively charged residues^21–23^. Finally, the unstructured C-terminal end of Hfq can also interact with certain RNAs to promote the exchange of RNAs^18, 21, 24^. The most refined model describing the interactions between Hfq and sRNAs/mRNAs has sorted sRNAs into two classes^25^. The proximal face of Hfq is generally important for binding of the sRNAs through their poly-U tail of the Rho-independent terminator. Class I sRNAs (such as RyhB and DsrA) use the rim as the second binding site, whereas class II sRNAs (such as ChiX and MgrR) use the distal face as the second binding site^25^. In addition, the preferred target mRNAs of the two classes of the sRNA are proposed to have the complementary binding sites on Hfq, *i.e.* class I sRNA-targeted mRNAs binding to the distal face, and class II sRNA-targeted mRNA binding to the rim, in order to efficiently form sRNA-mRNA complexes^25^. As for many other RBPs, the functions of Hfq are facilitated by its interactions with other essential protein factors. Particularly, RNase E, the key ribonuclease for processing and turnover of ribosomal RNA (rRNAs) and mRNAs, is known to interact with Hfq through its C-terminal scaffold region^26–28^. The Hfq-RNase E interactions can promote degradation of the sRNA-targeted mRNA^28–31^.

**Figure 1.**
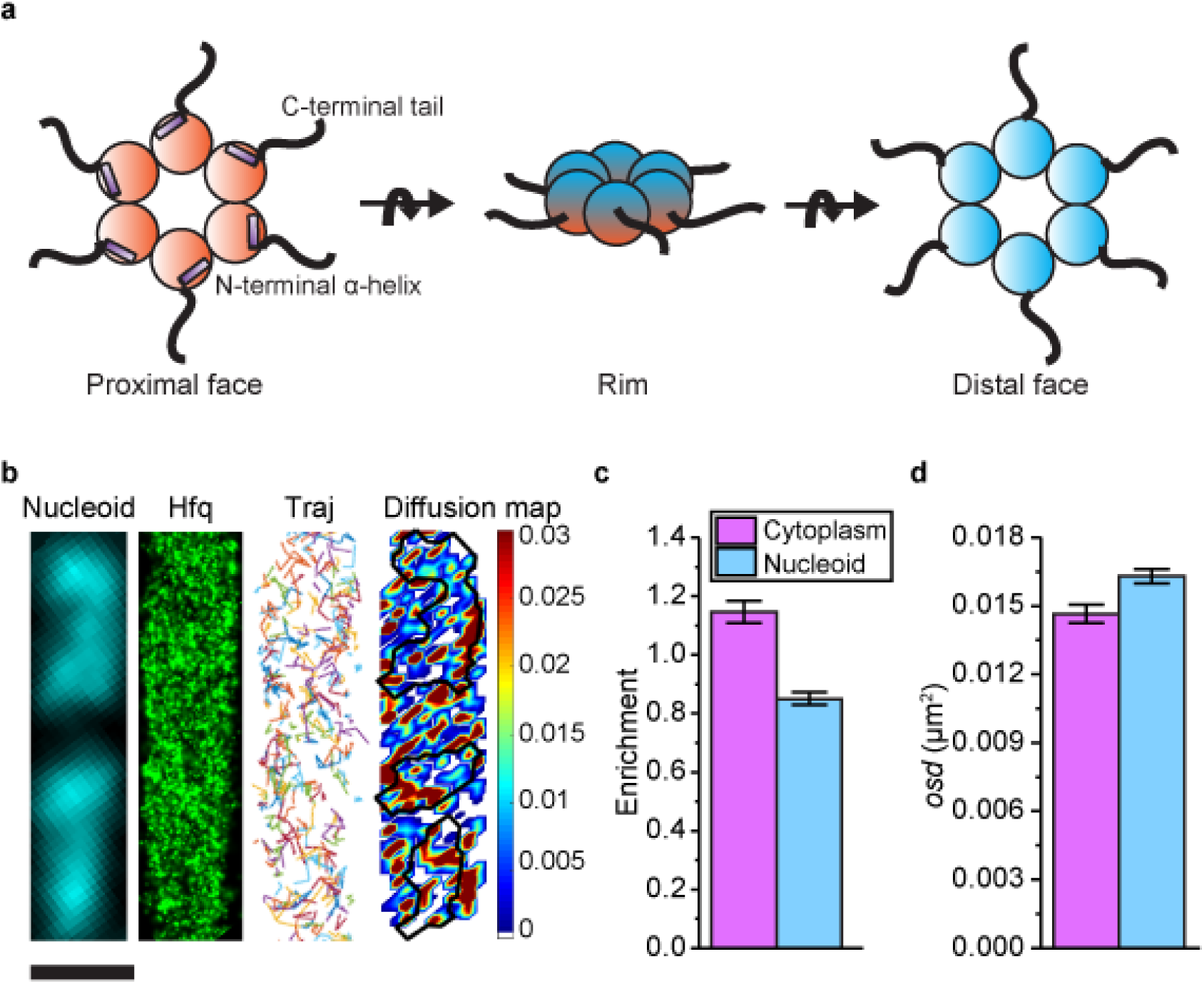
Diffusion and localization of Hfq during exponential growth. **a**, Schematic representation of Hfq with three RNA binding faces indicated. **b**, Representative image of WT hfq-mMaple3 in WT *rne* background during exponential growth under no treatment (NT) condition. Nucleoid is stained with Hoechst in live cells. 2D reconstruction image of Hfq-mMaple is shown in the black background. Different diffusion trajectories from tracking algorithm are shown in different colors (“Traj”). *osd* (unit: µm^2^) at each coordinate of the cell is shown as heatmap, in which the black curves show the nucleoid region defined by Hoechst staining. The scale bar represents 1 µm. **c**, Enrichment of Hfq localization is calculated for nucleoid region and cytoplasm region under NT condition. **d**, Average *osd* of Hfq within the nucleoid and cytoplasm regions under NT condition. Error bars in all plots represent the standard deviation (s.d.) from 2 experimental replicates, each containing ∼60 cells.

While Hfq is an abundant RBP in bacterial cells^32, 33^, it is still considered to be limiting, given the abundance of cellular mRNAs and sRNAs. Particularly, *in vitro* studies on specific sRNAs demonstrate that Hfq binds RNAs tightly with a dissociation constant of nM, and the Hfq-RNA complexes are stable with a lifetime of >100 minutes^34–36^. However, under stress conditions, induced sRNAs can regulate target mRNAs within minutes, raising a long-standing question of how sRNAs can rapidly access Hfq that might be tightly bound by pre-existing cellular RNAs. To address this question, a model of RNA exchange on Hfq, *i.e.* a RNA can actively displace another RNA from Hfq, was proposed to account for the fast sRNA-mediated stress response^2, 3, 37^. While *in vitro* biophysical experiments can be used to measure the affinity of RBP binding to many different RNAs under many different controllable conditions, the *in vitro* nature of these experiments makes it difficult to replicate the concentrations, compartmentalization, crowding, competitive binding and changes in cellular conditions that can affect the behavior and function of RBPs in live cells. Therefore, the mechanism(s) that can recycle Hfq from pre-bound RNAs in live cells remains to be elucidated.

In order to address this question in a cellular context, we sought to measure the diffusivity of Hfq in live *Escherichia. coli* cells, using single-molecule localization microscopy (SMLM)^38^, with a rationale that the diffusivity is affected by the molecular weight of the molecules, and therefore can report the interactions between Hfq with different cellular components. By measuring Hfq diffusivity under a variety of cellular conditions in combination with other biochemical assays, we demonstrate that Hfq dynamically changes its interactions with different RNAs in response to changes in cellular conditions, reveal a new sRNA-independent pathway for Hfq-regulated mRNA turnover, and illustrate that the two classes of sRNA can gain access to mRNA pre-bound Hfq through different mechanisms.

## Results

### Cellular Hfq freely diffuses in the absence of stress

Hfq was tagged by a photo-switchable fluorescent protein (FP), mMaple3^39^, at the C-terminus and the fused *hfq* gene was integrated into the genomic locus to replace the wild-type (WT) *hfq* (denoted as “*hfq-mMaple3*”, Methods). The strain harboring *hfq-mMaple3* showed comparable growth curve as the WT strain, whereas Δ*hfq* showed a growth defect (Figure S1). In addition, Hfq-mMaple3 showed activity comparable to WT Hfq protein, as revealed by Northern blots of RyhB-mediated *sodB* mRNA degradation, and MicA-mediated *ompA* mRNA degradation (Figure S2).

We performed single-particle tracking using SMLM in two dimensions (2D). Imaging conditions and parameters for applying tracking algorithm were optimized using fixed samples as the control (Figure S3). We first tracked Hfq-mMaple3 in live cells grown at exponential phase (referred as “<u>n</u>o treatment”, or “NT” case). In the NT condition, Hfq-mMaple3 exhibited a relatively uniform distribution within the cell (Figure 1b), consistent with the distribution revealed by the earlier live-cell imaging with Hfq tagged by a different FP (Dendra2)^40^. Quantification of Hfq-mMaple3 localization with DNA stained by Hoechst revealed a slightly higher cytoplasm enrichment than nucleoid localization in the NT condition (Figure 1c). We did not observe a helical organization along the longitudinal direction of the bacterial cell^41^, membrane localization^42^, or cell pole localization^43^, as reported in a few fixed-cell experiments. In addition, we calculated the one-step squared displacement (*osd*) of individual Hfq-mMaple3 protein at each time step and plotted the *osd* as a function of the cellular coordinate in a diffusivity heatmap (Figure 1b). The heatmap and quantification of the average *osd* suggest that Hfq diffuses similarly within the nucleoid and cytoplasmic region (Figure 1d).

### Binding of mRNAs to Hfq decreases its diffusivity primarily through the distal face of Hfq

We first tested the effect of mRNA on Hfq-mMaple3 diffusivity by treating the cells with rifampicin (Figure 2a), an antibiotic that inhibits transcription and results in the loss of most cellular mRNAs. We estimated the effective diffusion coefficient (D) by fitting the power function to the mean squared displacement (MSD) as a function of time lag (Δt) (Figure 2b). Transcription inhibition increased the diffusivity of Hfq-mMaple3 (Figure 2b), suggesting that a fraction of Hfq-mMaple3 proteins are associated with cellular mRNAs, consistent with previous reports^40^. As a control, mMaple3 protein alone did not show any changes in diffusivity upon rifampicin treatment (Figure S4).

**Figure 2.**
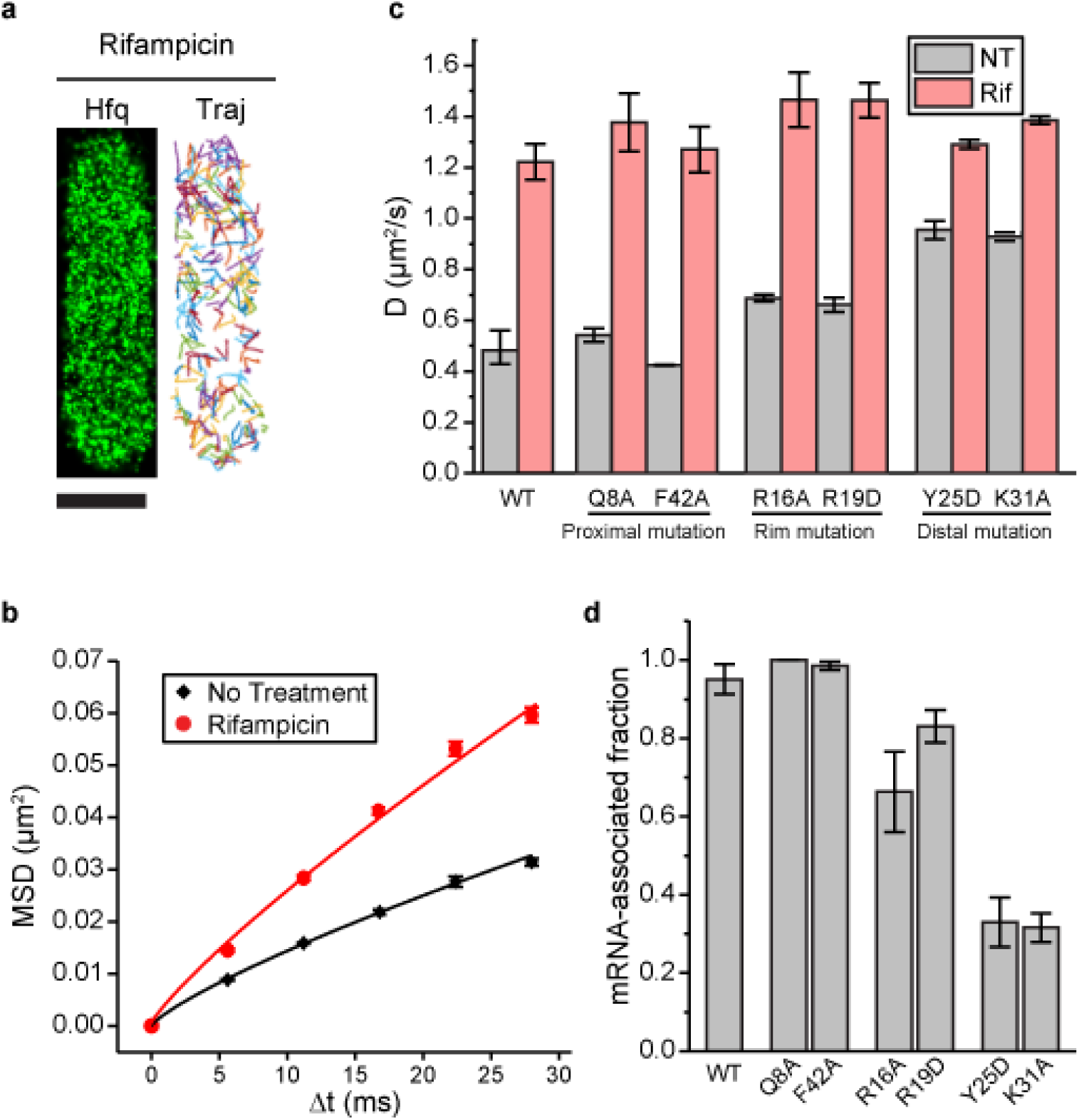
Binding of mRNAs to Hfq decreases its diffusivity primarily through the distal face of Hfq. **a**, Representative image of Hfq-mMaple3 with rifampicin treatment (Rif). 2D reconstruction image is shown in the black background (left), and different diffusion trajectories from tracking algorithm are shown in different colors (right). The scale bar represents 1 µm. **b**, Mean square displacement (MSD) is plotted against the time interval (Δt) for Hfq-mMaple3. The power law fitting curves are shown. **c**, Ensemble diffusion coefficients are plotted for WT and six mutations of Hfq-mMaple3 under NT and Rif conditions. **d**, mRNA-associated fraction for WT and mutant Hfq under NT case. Error bars in all plots represent the s.d. from 2 or 3 experimental replicates, each containing ∼100 cells.

We next introduced point mutations at each RNA binding face of Hfq-mMaple3^25, 44^, and imaged these mutant Hfq-mMaple3 proteins under NT and rifampicin treated conditions. With rifampicin treatment, all Hfq mutants exhibited similar diffusivity. However, mutations on different interfaces changed Hfq diffusivity in the NT case to different levels, suggesting that mutation on different faces changed the ability of Hfq to bind cellular mRNAs. Specially, both proximal face mutants (Q8A and F42A) exhibited similar diffusivity as the WT Hfq-mMaple3; both rim mutations (R16A and R19D) had a minor increase in diffusivity under NT condition; and both distal face mutations (Y25D and K31A) led to a large increase in the diffusivity under NT condition (Figure 2c). Comparison of the WT and the mutant Hfq-mMaple3 proteins supports conclusions that Hfq binds mRNAs in the cell and that binding of mRNAs is primarily achieved through the interactions with the distal face of Hfq, whereas the rim also contributes to the mRNA binding in a minor way.

### Most Hfq proteins are occupied by mRNAs in the cell during exponential growth

Under our experimental conditions, mMaple3 protein alone had a D of 1.8±0.2 µm^2^/s (fit with a power function) or 2.7±0.2 µm^2^/s (fit with a linear function) (Figure S4), close to the reported range of D for fluorescent proteins (∼27 kDa) (3-8 µm^2^/s)^45^. WT and mutant Hfq-mMaple3 proteins upon rifampicin treatment consistently showed a diffusion coefficient of 1.2-1.5 µm^2^/s (Figure 2c). Considering the power-law dependence of D on Mw of biomolecules^45^, such a change in D corresponds to a 2-4 fold change in Mw (∼50-100 kDa), which is smaller than hexameric Hfq-mMaple3 (∼220 kDa). This observation suggests that in the absence of RNA binding, a fraction of Hfq may exist as monomer in the cell. WT Hfq-mMaple3 in the NT case had a D of 0.50±0.07 µm^2^/s, corresponding to a Mw of 2.1 MDa, again assuming the power-law relationship between D and Mw. Considering the average length of bacterial mRNAs to be 1 kb (∼330 kDa), and Mw of bacterial ribosome (∼2.5 MDa), this reduction in D supports the interpretation that a significant fraction of WT Hfq proteins are associated with mRNAs in the NT case, and that a fraction of the associated mRNAs are translated by the ribosomes. In addition, previous experiments have measured the diffusion coefficient of ribosomes or ribosomal complexes to be 0.04-0.5 µm^2^/s^46–48^. The average D of WT Hfq-mMaple3 under NT condition is at the upper limit of range for ribosomes, again indicating the D value of mRNA-associated Hfq very likely represents an average value of both untranslated and translated mRNAs.

In order to estimate the fraction of mRNA-associated Hfq-mMaple3, we plotted log(*osd*) in a histogram. Consistent with D values, distribution of *osd* overall shifted to larger values with rifampicin treatment compared to the NT case (Figure S5). We fit the distribution of log(*osd*) of the rifampicin treated case with single Gaussian peak and used the fit parameters (the center and the width) to constrain the fitting for other Hfq-mMaple3 constructs under different conditions. For the WT Hfq-mMaple3, in the NT case, about 95±4% of the population was mRNA-associated, consistent with previous hypothesis that Hfq proteins are largely occupied in the cell^2, 3, 37^. Y25D and K31A mutants had the most compromised mRNA binding ability, with 33±6% and 32±4% of the population was mRNA-associated, respectively (Figure 2d).

### Hfq is deficient in releasing mRNAs without interactions with RNase E

Hfq has been demonstrated to interact with the C-terminal scaffold region of RNase E^26–28^. To study whether the interaction with RNase E affects the diffusivity of Hfq, we imaged Hfq-mMaple3 in two RNase E mutant strains. The *rne131* mutant strain has RNase E truncated by the last 477 amino acid residues^49^, therefore, while it maintains its nuclease activity, this mutant cannot interact with Hfq. The *rne*Δ*14* mutant has a smaller fraction of the C-terminal scaffold (residues 636-845) deleted, encompassing the Hfq binding region^50^. In both RNase E mutant backgrounds, the diffusivity of Hfq-mMaple3 became less sensitive to transcription inhibition by rifampicin compared to the WT *rne* case (Figure 3a and 2c). 40-50% of Hfq-mMaple3 remained mRNA-associated upon rifampicin treatment in the RNase E mutant backgrounds (Figure 3b). These observations suggest that without the Hfq-RNase E interaction, more mRNAs remain bound to Hfq, indicating that Hfq may help deliver the associated mRNA to RNase E for degradation.

**Figure 3.**
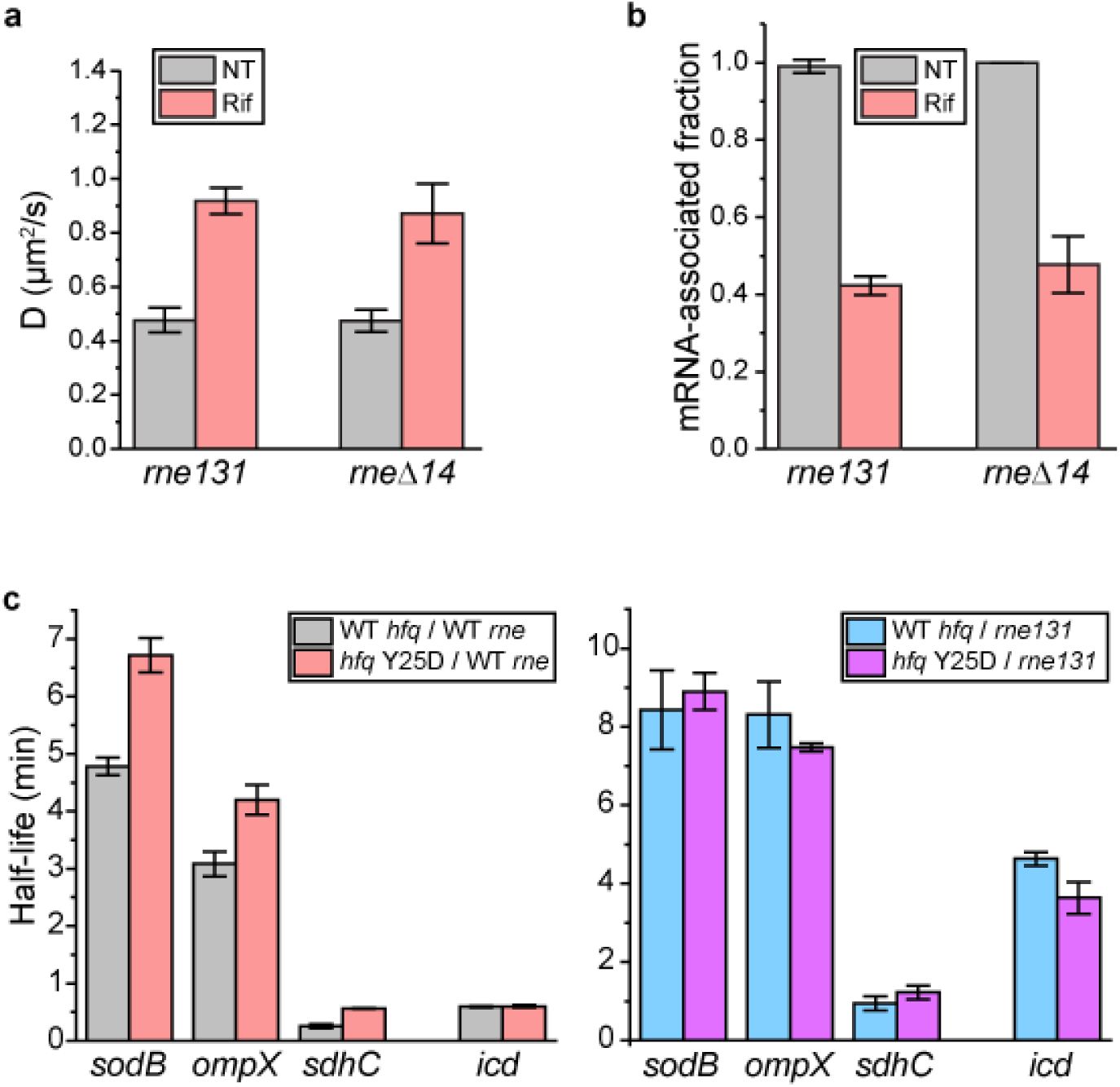
Hfq-RNase E interaction contributes to the degradation of mRNAs. **a**, Diffusion coefficients are plotted for WT hfq-mMaple3 in the *rne131* and *rne*Δ*14* backgrounds under NT and Rif conditions. **b**, mRNA-associated fraction of WT Hfq in the *rne131* and *rne*Δ*14* backgrounds under NT and Rif conditions. **c**, Half-lives of selected mRNAs determined by Northern blots (Figure S6). sRNA regulators of these mRNAs were knocked out (Δ*ryhB*Δ*fnrS* for *sodB*, Δ*cyaR*Δ*micA* for *ompX* and Δ*ryhB*Δ*spf*Δ*rybB* for *sdhC*). Error bars in all plots represent the s.d. from 2 or 3 experimental replicates. Each imaging experiment contains ∼100 cells.

### Hfq-RNase E interaction contributes to the degradation of Hfq-associated mRNAs

To further test the hypothesis that Hfq plays a role in the turnover of certain mRNAs, we used Northern blots to measure the half-lives of selected mRNAs that are known to interact with Hfq in four backgrounds: (1) WT *hfq-mMaple3* + WT *rne*, (2) WT *hfq-mMaple3* + *rne131* mutant, (3) *hfq* Y25D*-mMaple3* + WT *rne*, (4) *hfq* Y25D*-mMaple3* + *rne131* mutant (Figures 3c and S6). In addition to the genetic background of *hfq-mMaple3* and *rne*, we also knocked out the corresponding sRNA regulators of the selected mRNAs (Δ*ryhB*Δ*fnrS* for *sodB*, Δ*cyaR*Δ*micA* for *ompX* and Δ*ryhB*Δ*spf*Δ*rybB* for *sdhC*). In the WT *rne* background, all tested mRNAs (except for *icd*, to be discussed later) showed an increased half-life in the *hfq* Y25D*-mMaple3* background compared to WT *hfq-mMaple3* (Figure 3c), suggesting that association with Hfq facilitates the turnover of these mRNAs. In the *rne131* mutant background, while all mRNAs showed an increased lifetime of 2 to 7-fold compared to the WT *rne* background, consistent with a compromised activity in the *rne131* mutant^49^, the difference in the mRNA half-lives between WT *hfq-mMaple3* and *hfq* Y25D*-mMaple3* backgrounds was largely diminished (Figure 3c). This result indicates that in the absence of Hfq-RNase E interaction, association with Hfq or not does not change the mRNA turnover. As a negative control, *icd* mRNA, which does not have a putative Hfq binding site, exhibited similar half-lives under WT *hfq* and h*fq* Y25D backgrounds (Figure 3c). These results collectively support that besides the sRNA-mediated pathway, Hfq can regulate certain mRNAs’ half-lives by bringing the mRNAs to RNase E for degradation.

### sRNAs can displace Hfq from mRNAs in a face-dependent manner

We next examined the effect of sRNAs on the diffusivity of Hfq-mMaple3. We induced expression of different sRNAs, including RyhB, a class I sRNA, ChiX a class II sRNA, and a sRNA that is less clearly defined between these two classes, SgrS^25^. Whereas overexpression of RyhB or SgrS did not cause any noticeable changes in the Hfq-mMaple3 diffusivity or mRNA-associated fraction, overexpression of ChiX dramatically increased its diffusivity and lowered the mRNA-associated fraction (Figure 4a and b).

**Figure 4.**
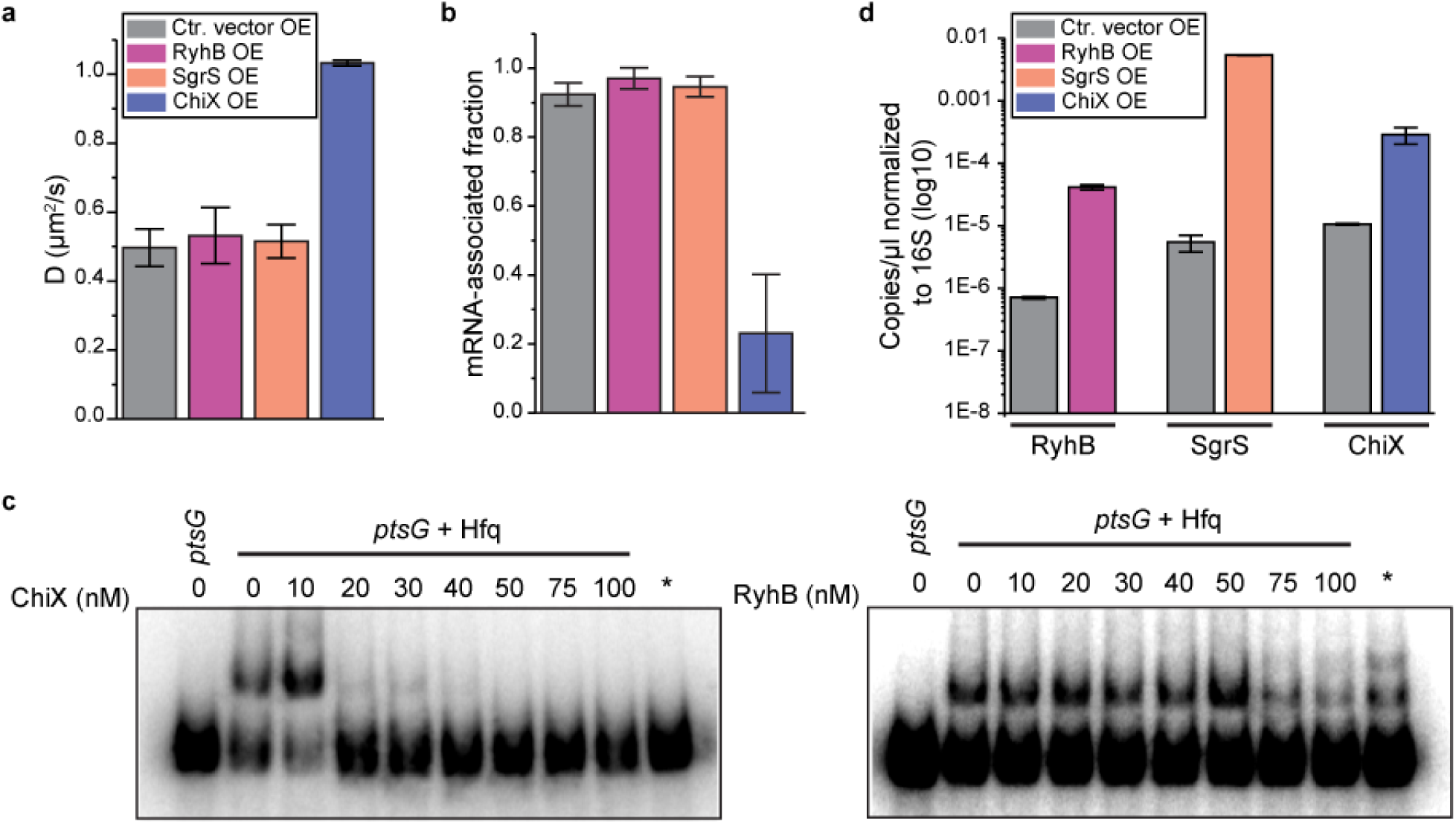
sRNAs can displace mRNA from in a face-dependent way. **a,** Diffusion coefficients of Hfq-mMaple3 with RyhB, SgrS or ChiX over-expressed from an IPTG inducible promoter. **b,** mRNA-associated fraction for Hfq, for the same cases as in a. Error bars in all plots represent the s.d. from 2 experimental replicates, each containing ∼100 cells. **c,** Competition of ChiX and RyhB on pre-occupied Hfq. 20 nM of a *ptsG* RNA fragment was pre-incubated with 100 nM Hfq before addition of increasing concentrations of ChiX or RyhB sRNA. (*) = Hfq (100 nM) and ChiX or RyhB (100 nM) were simultaneously added to 20 nM *ptsG* fragment. Data is representative of 3 independent experiments. K_d_ measurements of RyhB, ChiX and ptsG fragment binding to Hfq are shown in Figure S7. **d**, Abundance of RyhB, SgrS or ChiX in a control condition (Ctr. vector) or for each sRNA over-expressed from an IPTG inducible promoter, normalized to 16S rRNA measured by ddPCR.

As described above, the distal face is the primary binding site for mRNAs in the cell (Figure 2c). Since ChiX requires binding at both the proximal and distal faces, we expect the diffusivity of Hfq to increase after shifting from mRNA-bound Hfq to ChiX-bound Hfq. Due to the relatively small molecular weight of sRNAs (∼50-300 nucleotides in length), sRNA-bound Hfq-mMaple3 has similar diffusivity as free Hfq-mMaple3. We then checked if ChiX could compete with mRNAs for binding to Hfq *in vitro* using electrophoretic mobility shift assay (EMSA). A radiolabeled fragment of *ptsG* mRNA was pre-incubated with purified Hfq protein and then chased with unlabeled ChiX. Consistent with the *in vivo* results, ChiX can effectively displace *ptsG* from Hfq (Figure 4a and c, left panel).

Overexpression of RyhB or SgrS did not cause any significant changes in the Hfq-mMaple3 diffusivity or the corresponding mRNA-associated fraction (Figure 4a and b). We reasoned that there might be two possibilities. First, since class I sRNAs bind through the proximal face and the rim of Hfq, it can bind to the mRNA-free Hfq or co-occupy the mRNA-bound Hfq to generate sRNA-bound Hfq or sRNA-mRNA-co-bound Hfq, respectively. Second, class I sRNAs cannot effectively compete against mRNAs for Hfq binding, therefore most Hfq proteins remain associated with mRNAs. To distinguish these two possibilities, we performed an EMSA competition assay using RyhB as an example. Results show that RyhB cannot displace the radiolabeled *ptsG* from Hfq, but rather generates an additional upper-shifted band compared to the band of *ptsG*-Hfq complex, supporting that RyhB and *ptsG* can co-occupy Hfq (Figure 4c, right panel). In addition, droplet digital PCR (ddPCR) performed in the same conditions as the diffusivity assays showed that RyhB level was comparable to ChiX (Figure 4d). Since RyhB stability is highly dependent on association with Hfq^51^, EMSA and ddPCR results suggest that both *in vitro* and *in vivo*, RyhB can effectively access mRNA-occupied Hfq through co-occupying Hfq from the proximal face.

To summarize, our results collectively suggest that both class I and class II sRNAs can access mRNA-occupied Hfq *in vivo*. Class I sRNAs can co-occupy the Hfq protein with an mRNA through different binding sites whereas class II sRNAs can directly compete against the mRNA at the distal face. Interestingly, fluorescence *in situ* hybridization (FISH) showed a much stronger signal for RyhB compared to ChiX (Figure S8), even though their levels are similar revealed by ddPCR (Figure 4d). The weaker hybridization signal for ChiX is very likely a reflection of the larger protected region by Hfq on both distal and proximal faces, hindering FISH probe binding.

### Class II sRNAs require interaction with the proximal face and a strong AAN motif to compete for Hfq binding

We next sought to understand the molecular features that make ChiX a strong competitor for Hfq binding. When overexpressed in the *hfq* Q8A-*mMaple3* background (proximal face mutation), ChiX lost its capability to displace mRNAs from the mutant Hfq (Figure 5b and c), suggesting that additional binding affinity provided by the proximal face of Hfq is critical for displacing other RNAs from the distal face. *E. coli* Hfq prefers a (A-A-N)_n_ sequence for distal face binding, where N can be any nucleotide, and each monomer binds to one A-A-N repeat^18^. ChiX contains four AAN motifs (Figure 5a). We next determined the effect of AAN motifs on conferring the competitive binding to Hfq over mRNAs. We generated and overexpressed ChiX mutants with one or two AAN motif(s) deleted (Figure 5a) and found that the fraction of remaining mRNA-bound Hfq increased when the number of AAN motifs decreased (Figure 5a and c). Notably, the levels of WT ChiX in the *hfq* Q8A-*mMaple3* background, and the ChiX mutants in the WT *hfq-mMaple3* background remained similar as the WT ChiX in WT *hfq-mMaple3* background (Figure 4d and 5d), confirming that the difference we observed in the mutant cases was not due to a change in the cellular ChiX level.

**Figure 5.**
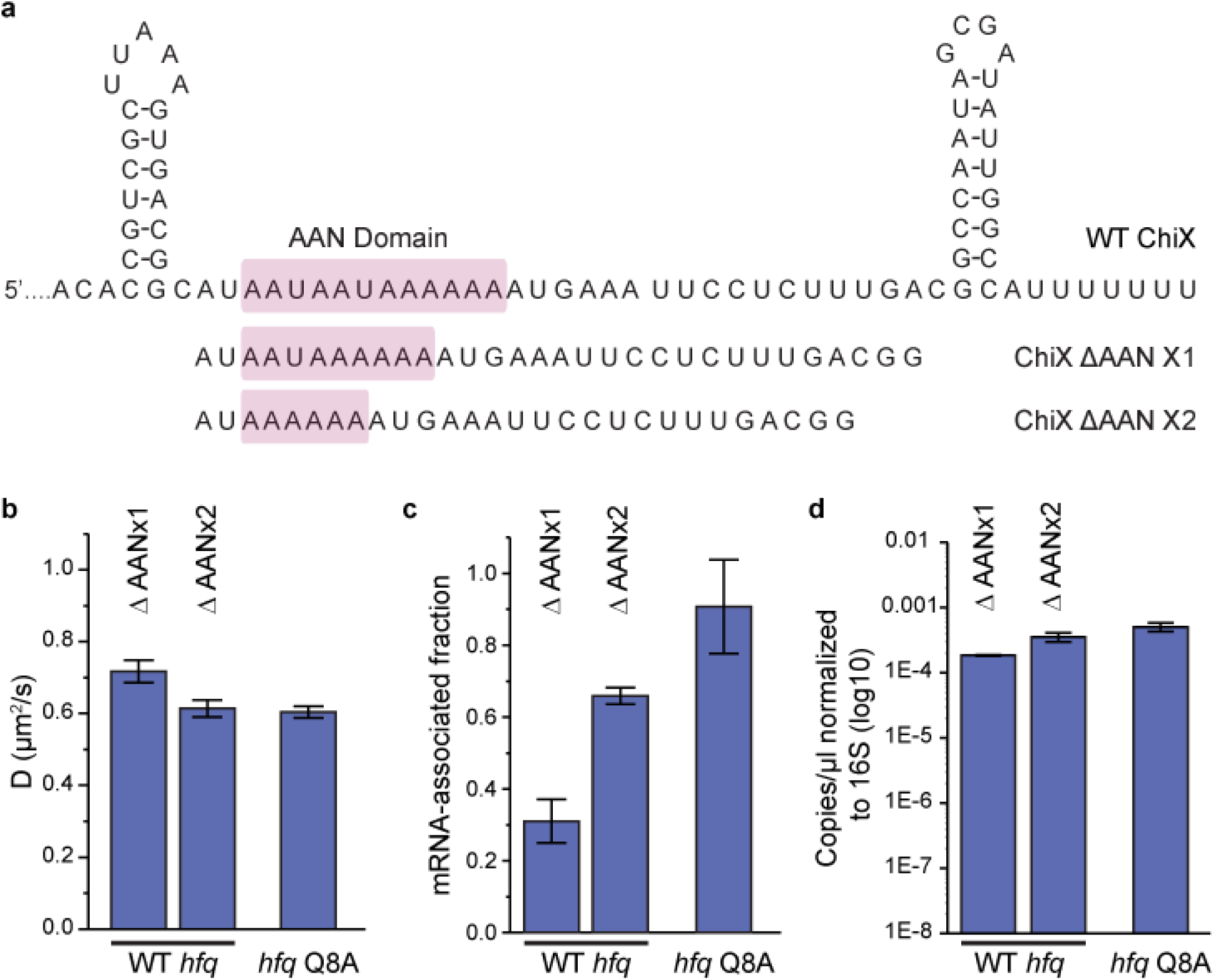
Both interactions at the proximal and distal face contributes to the competitive binding of ChiX. **a,** Sequences of WT ChiX and two ChiX mutants (with one or two AAN motif deleted). **b,** Diffusion coefficients and **c,** mRNA-associated fraction of Hfq-mMaple3 with ChiX mutants over-expressed, or Hfq Q8A-mMaple with WT ChiX over-expressed. **d,** Abundance of ChiX ΔAANx1-2 mutants in the WT *hfq-mMaple3* and WT ChiX in the *hfq* Q8A*-mMaple3* background normalized to 16S rRNA measured by ddPCR.

## Discussion

Using single-particle tracking, we resolved different diffusivity states of Hfq proteins in live cells, reporting on the interactions with different cellular RNAs. Specifically, free Hfq and sRNA-bound Hfq proteins have the highest diffusivity, and association of mRNAs reduces the diffusivity of Hfq. Utilizing the different diffusivities of mRNA-associated Hfq and mRNA-free Hfq, we directly observed the transition of major interaction partners of Hfq in response to changes in cellular conditions. During exponential growth when sRNAs are least expressed, Hfq proteins are largely occupied by mRNAs through the distal face. Consistent with *in vitro* binding studies^34–36^, intracellular interactions between Hfq and the mRNA are fairly stable, as indicated by the observation that Hfq cannot release the bound mRNA efficiently in the absence of the interaction with RNase E upon rifampicin treatment. On the other hand, the mRNA binding and RNase E binding capability of Hfq allows it to serve as a “shuttle” by bringing mRNA to RNase E to promote turnover of certain Hfq-associated mRNAs in a sRNA-independent manner (Figure 6a). Therefore, Hfq-RNase E interaction not only regulates the stability of certain Hfq-associated mRNAs, but also provides one mechanism to recycle Hfq from these mRNAs.

**Figure 6.**
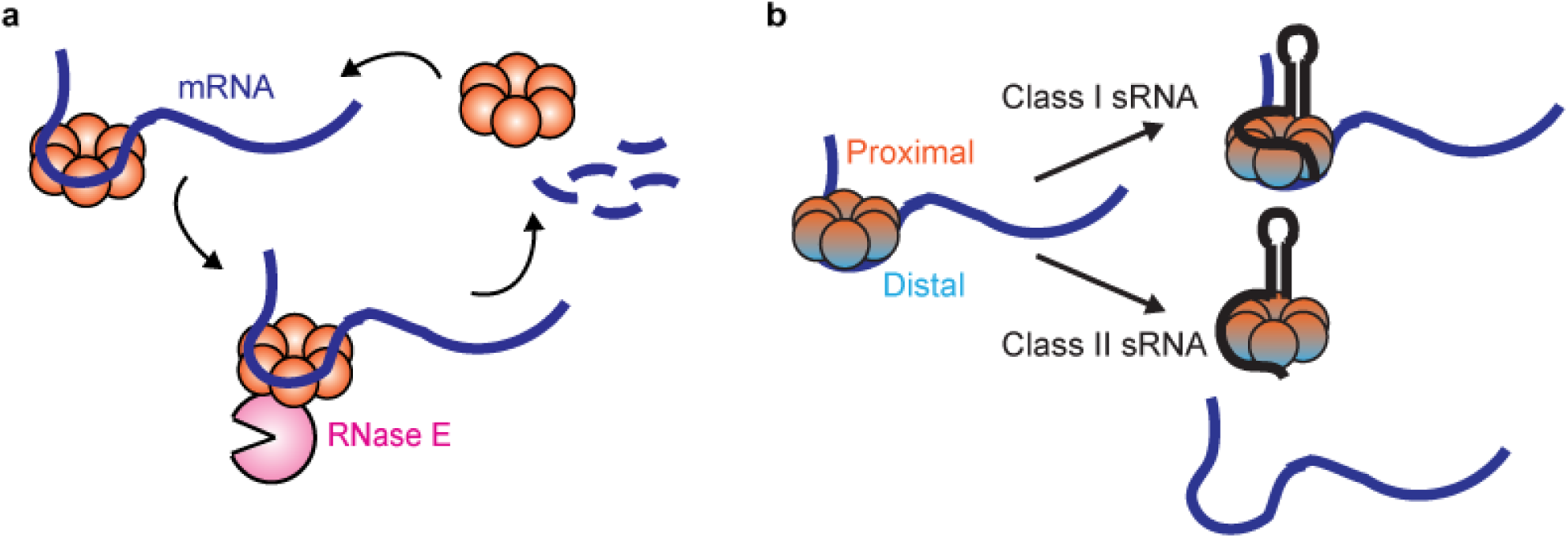
Dynamic interactions between Hfq and cellular RNAs. **a,** Hfq facilitates the degradation of certain Hfq-bound mRNAs through the recruitment of RNase E. **b,** Class I or Class II sRNAs can get access of mRNA-associated Hfq through co-occupying different binding sites of Hfq simultaneously, or displacing mRNA from the distal face of Hfq respectively.

Our results suggest that for the selected mRNAs, Hfq can regulate their turnover through a direct recruitment of RNase E. Mechanisms of sRNA-independent Hfq-mediated regulation on mRNA turnover have been reported. First, binding Hfq, or Hfq in complex with other proteins such as Crc, at the ribosome binding site of the mRNA can repress translation^6–8^, therefore indirectly increasing the mRNA degradation due to de-protection of the translating ribosomes against RNase E. Since this translation-dependent regulation of Hfq does not require Hfq-RNase E interaction, if this mechanism applied to the mRNAs we tested, we would expect the difference in half-life between the cases of WT *hfq* and *hfq* Y25D to be similar in the WT *rne* and *rne131* backgrounds. The observation that the difference in half-life due to Hfq binding is eliminated in the *rne131* background suggests the turnover of these mRNAs is not through regulation at the translation level. Second, binding Hfq may recruit polyA polymerase (PAP) and polynucleotide phosphorylase (PNPase) to stimulate polyadenylation at the 3’ end, and therefore promote degradation^9, 10^, an action that may also require interactions with the C-terminal scaffold region of RNase E. However, this mechanism also cannot fully explain our results for three reasons: (1) Previous studies suggested that Hfq-stimulated polyadenylation prefers Hfq binding at the 3’ termini of mRNAs containing Rho-independent transcription terminator^10^. However, in our selected mRNAs, they do not all utilize Rho-independent termination. (2) The Y25D mutant is more likely to hinder mRNA binding through AAN motif on the distal face, rather than through polyU stretch in the Rho-independent terminator, assuming binding of Hfq to the Rho-independent terminator would most likely be through interactions of between the polyU tail and the proximal face. (3) Our imaging results on Hfq mutants suggest that Hfq binds mRNA mostly through the distal face, indicating that the population of mRNAs with Hfq binding at a Rho-independent terminator would only be a minor fraction, highly consistent with the CLIP-seq results of Hfq^14^. Therefore, it is more likely that the regulation for the selected mRNAs is through direct recruitment of RNase E rather than through Hfq-stimulated polyadenylation mechanism. Nevertheless, Hfq can potentially regulate mRNA turnover through a combination of these three mechanisms in a gene-specific manner.

Under the conditions when a specific sRNA is highly induced, we demonstrate how the sRNA can quickly gain access to mRNA pre-occupied Hfq proteins (Figure 6b). Our results are reminiscent of a previously proposed model of Hfq interacting with sRNAs and mRNAs in a face-dependent manner^25^. While the distal face of Hfq is the primary binding site for cellular mRNAs, the rim has a minor binding role, suggesting that the majority of the Hfq-bound mRNAs are class I mRNAs, and a minority are class II mRNAs. This observation is consistent with the previous findings that most of the sRNAs are class I sRNAs^25^. Both classes of sRNAs can easily access Hfq upon induction, albeit with different mechanisms. Surprisingly, class I sRNAs can directly co-occupy Hfq through the binding faces that are non-overlapping with the class I mRNA binding face, without the need to displace the pre-occupied mRNA. In contrast, class II sRNAs, such as ChiX, can either co-occupy the minor fraction of Hfq associated with the class II mRNA at the rim, or more likely, can effectively displace class I mRNAs from the distal face. In both cases, the mRNA-occupied Hfq proteins are in standby mode for sRNA binding if needed. The displacement of mRNA by the class II sRNA requires both the interactions at the proximal face of Hfq and higher AAN motif number to outcompete mRNAs for the binding at the distal face. In addition, we propose that the competitive binding by the class II sRNA is likely to occur stepwise, with binding at the proximal face happening first, followed by the displacement of mRNA form the distal face, which is supported by the observation that with the Hfq proximal face mutation, ChiX cannot displace mRNAs, even with a strong AAN motif. Importantly, our mutational work demonstrates that it should be possible to engineer Hfq-RNA interactions *in vivo*. Given the complete absence of sRNA-mediated gene regulation in eukaryotes, our results provide a framework for developing biotechnological tools that will potentially enable precise control over the sequestration, degradation, and/or expression of target mRNAs in eukaryotes.

## Methods

### Bacterial Strains

Transfer of the Linker-mMaple3-Kan sequence at the 3’ end of chromosomal *hfq* gene was achieved by following the PCR-based method^52^ with a few modifications. First, a PCR (PCR1) was performed using plasmid pZEA93M as template to amplify the mMaple3 sequence (oligos EM4314-4293). Then, to add sequence homology of *hfq* gene, a second PCR (PCR2) was performed using PCR1 as template (oligos EM4313-4293). The final PCR product (PCR3), containing a flippase recognition target (FRT)-flanked kanamycin resistance cassette was generated from pKD4 plasmid with primers PCR2 and EM1690 carrying extensions homologous to the *hfq* gene. PCR3 was then purified and transformed to WT (EM1055), *hfq* Y25D (KK2562), *hfq* Q8A (KK2560), *hfq* K31A (AZZ41) or *hfq* R16A (KK2561) strains containing the pKD46 plasmid using electroporation, to obtain strains with *hfq*-Linker-mMaple3-Kan, *hfq* Y25D-Linker-mMaple3-Kan, *hfq* Q8A-Linker-mMaple3-Kan, *hfq* K31A-Linker-mMaple3-Kan, and *hfq* R16A-Linker-mMaple3-Kan, respectively. Hfq mutations F42A and R19D were obtained by performing PCRs on fusion strains KP1867 (Hfq-linker-mMaple3) with oligos EM4704-1690 (Hfq F42A) or EM4705-1690 (Hfq R19D). Fragments were then transformed to WT (EM1055) containing the pKD46 plasmid, following induction of the λ Red. P1 transduction was used to transfer the linked fluorescent protein and the antibiotic resistance gene into a WT (EM1055), *rne*131 (EM1377) or *rne*Δ14 (EM1376) strains.

*fnrS*, *micA* and *spf* knock-outs were obtained through transformation of PCR products into EM1237 after induction of λ *red* and selecting for kanamycin or chloramphenicol resistance. P1 transduction was used to transfer the knock-out mutations and the antibiotic resistance gene into appropriate strains. Selection was achieved with kanamycin or chloramphenicol. When necessary, FRT-flanked antibiotic resistance cassettes were eliminated after transformation with pCP20, as described^52^. All constructs were verified by sequencing and are listed in Table S1. Oligonucleotides used for generating constructs are listed in Table S2.

### Plasmids

*E. coli* MG1655 sRNA genes *sgrS*, *ryhB,* and *chiX* were inserted into the pET15b low copy number vector plasmid (kind gift from Perozo lab) to create plasmids pET16b-RyhB, pET15b-SgrS and pET15b-ChiX using Gibson Assembly (in house) using oligos listed in Table S2. The *chiX ΔAAN* mutants were made using site directed mutagenesis. Primers EH159, EH160 and EH161 homologous to *chiX* while excluding the AAN domain were used to amplify the plasmid. The products were phosphorylated (NEB M0201S) and ligated (NEB M0202S) before transformation.

Cloning of pBAD-*micA* was performed by PCR amplification of *micA* (oligos EM2651-2652) on WT strain (EM1055). The PCR product was digested with SphI and cloned into a pNM12 vector digested with MscI and SphI. All constructs were verified by sequencing and are listed in Table S1. Oligos used for generating the constructs are listed in Table S2.

### Growth conditions for imaging experiments

Overnight cultures of *E. coli* strains were diluted by 1:100 in MOPS EZ rich defined medium (Teknova). 0.2% glucose was used as the carbon source for imaging Hfq-mMaple3 WT and mutants under NT and rifampicin treated conditions. 0.2% glycerol was used as the carbon source with 100 µg/mL ampicillin for mMaple3 control (SP191). 0.2% fructose was used as the carbon source with 100 µg/mL ampicillin for cases with sRNA overexpression. Cultures were grown at 37 °C aerobically. Plasmid-encoded sRNAs and plasmid-encoded mMaple3 proteins were induced by IPTG (1 mM and 100∼400 μM respectively) when the OD_600_ of the cell culture was ∼0.1. Induced cells were grown for ∼45 minutes before imaging. For the rifampicin treatment, rifampicin was added to a final concentration of 200 μg/mL when the OD_600_ of the cell culture was ∼0.2, and the cells were incubated for 15 minutes before imaging.

### Growth curve measurement

The bacterial strains were grown overnight in LB or MOPS EZ-rich medium containing 0.2% glucose. Cultures were diluted to 6 × 10^6^ cells/mL in their respective medium and samples were prepared in triplicate by mixing 50 µL of cells and 50 µL of fresh media to obtain 3 × 10^6^ cells/mL. Assay were performed in Microtest plate, 96-well, flat base, polystyrene, sterile (Sarstedt) and growth was monitored using Epoch 2 Microplate Spectrophotometer reader (BioTek) with the following settings: OD= 600 nm, Temperature= 37 °C, Reading= every 10 min for 22 h, Continuous shaking.

### RNA extraction and Northern blot analysis

Total RNA was extracted following the hot-phenol protocol as described^53^. To test the function of the mMaple3-tagged Hfq and compare that with the WT Hfq, cells were grown in LB to the OD_600_ of 0.5 and either RyhB was induced by adding of 2.2′-dipyridyl in a WT *hfq* or in an *hfq*-*mMaple3* background, or MicA was induced by addition of 0.1% arabinose (ara) in a Δ*micA* WT *hfq* or in a Δ*micA hfq*-*mMaple3* background (pBAD-*micA*).

Determination of RNA half-life was performed in MOPS EZ rich defined medium (Teknova) with 0.2% glucose by addition of 500 μg/mL rifampicin to the culture at the OD_600_ of 0.5 before total RNA extraction. Northern blots were performed as described previously^54^ with some modifications. Following total RNA extraction, 5∼10 μg of total RNA was loaded on polyacrylamide gel (5% acrylamide 29:1, 8 M urea) and 20 μg was loaded on agarose gel (1%, 1X MOPS). Radiolabeled DNA and RNA probes used in this study are described in Table S2. The radiolabeled RNA probes used for Northern blot analysis were transcribed with T7 RNA polymerase from a PCR product to generate the antisense transcript of the gene of interest^55^. Membranes were then exposed to phosphor storage screens and analyzed using a Typhoon Trio (GE Healthcare) instrument. Quantification was performed using the Image studio lite software (LI-COR).

The decay rate of mRNA degradation was calculated as previously described^56^. Briefly, the intensity of Northern blot at each time point upon adding rifampicin was normalized to the intensity at time zero, and was fit by a piecewise function in the log space:

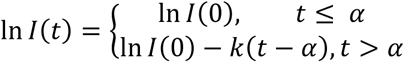

Where *I*(*t*) is the normalized intensity at time t, *I*(0) is the normalized intensity at time zero, *k* is the rate of exponential decay and *α* is the duration of the initial delay before the exponential decay begins. The reported half-lives (*τ*) is calculated by *τ* = *log* (2)/*k*.

### Droplet Digital PCR

Droplet Digital PCR (ddPCR) was performed on total RNA extracted following the hot-phenol protocol^53^ from cells grown in MOPS EZ rich defined medium containing 0.2% fructose (Teknova) with 50 µg/mL ampicillin. 1mM IPTG was added at OD_600_=0.1 for 1 h before total RNA extraction. Samples were treated with 8 U Turbo DNase (Ambion) for 1 h. RNA integrity was assessed with an Agilent 2100 Bioanalyzer (Agilent Technologies). Reverse transcription was performed on 1.5 µg total RNA with Transcriptor reverse transcriptase, random hexamers, dNTPs (Roche Diagnostics), and 10 U of RNase OUT (Invitrogen) following the manufacturer’s protocol in a total volume of 10 µL.

Droplet Digital PCR (ddPCR) reactions were composed of 10 µL of 2X QX200 ddPCR EvaGreen Supermix (Bio-Rad), 10 ng (3 µL) cDNA,100 nM final (2 µL) primer pair solutions and 5 µL molecular grade sterile water (Wisent) for a 20 µL total reaction. Primers are listed in Table S2. Each reaction mix (20 µL) was converted to droplets with the QX200 droplet generator (Bio-Rad). Droplet-partitioned samples were then transferred to a 96-well plate, sealed and cycled in a C1000 deep well Thermocycler (Bio-Rad) under the following cycling protocol: 95 °C for 5 min (DNA polymerase activation), followed by 50 cycles of 95 °C for 30 s (denaturation), 59 °C for 1 min (annealing) and 72 °C for 30 s (extension) followed by post-cycling steps of 4 °C for 5 min and 90 °C for 5 min (Signal stabilization) and an infinite 12 °C hold. The cycled plate was then transferred and read using the QX200 reader (Bio-Rad) either the same or the following day post-cycling. The concentration reported is copies/μL of the final 1x ddPCR reaction (using QuantaSoft software from Bio-Rad)^57^.

### Hfq purification

Hfq was purified following the previously described procedure^58^ with modifications. Briefly, strain EM1392 containing pET21b-*hfq* was grown at 37 °C in LB medium supplemented with 50 μg/mL ampicillin and 30 μg/mL chloramphenicol until it reached an OD_600_=0.6. Hfq expression was induced by addition of 5 mM IPTG (Bioshop) for 3 h. Cells were pelleted by centrifugation (15 min, 3825*g*) and resuspended in 4 mL Buffer C (50 mM Tris-HCl pH 7.5, 1 mM EDTA, 50 mM NH_4_Cl, 5% glycerol)^59^ supplemented with 30 U Turbo DNase (Ambion). Cells were lysed by sonication for 4 min (amplitude 25%, cycles of 5 sec sonication, 5 sec on ice) and samples were cleared by centrifugation (45 min, 12,000*g*). The supernatant was incubated at 80 °C for 10 min, centrifuged again (20 min, 12,000*g*) and cleared by filtration.

The protein extract was loaded onto a 1 mL HiTRAP Heparin column HP (GE Healthcare Life Sciences, 17-0406-01) equilibrated with Buffer A (50 mM Tris-HCl pH 8.0, 50 mM NaCl, 50 mM KCl, 1 mM EDTA, 5% glycerol). After washes, the protein was eluted with a linear NaCl gradient (0.05 M – 1 M NaCl) in Buffer A. Fraction samples were loaded on SDS-PAGE and stained with Coomassie-Blue. Hfq-containing fractions were dialyzed against a dialysis buffer (50 mM Tris-Cl pH 7.5, 1 mM EDTA pH 8.0, 5% Glycerol, 0.25 M NH_4_Cl). Glycerol concentration was brought up to 10% and protein content was quantified by BCA assay (Thermo Scientific™).

### EMSA

DNA templates containing a T7 promoter were synthesized by PCR amplification on genomic DNA using oligonucleotides EM88-EM1978 (T7-*ryhB*), T7-ChiX(F)-T7-ChiX(R) (T7-*chiX*) or T7-ptsG(F)-T7-ptsG(R) (T7-*ptsG*). Briefly, templates were incubated for 4 h at 37 °C in RNA Transcription Buffer (80 mM HEPES-KOH pH 7.5, 24 mM MgCl_2_, 40 mM DTT, 2 mM spermidine) in the presence of 5 mM NTP, 40 U porcine RNase Inhibitor (in house), 1 μg pyrophosphatase (Roche) and 10 μg purified T7 RNA polymerase (in house). Samples were treated with 2 U Turbo DNase (Ambion) and purified on polyacrylamide gel (6% acrylamide:bisacrylamide 19:1, 8 M urea). When necessary, transcripts were dephosphorylated using 10 U Calf Intestinal Phosphatase (NEB) and were 5’ end-radiolabeled with [Ɣ-^32^P]-ATP using 10 U T4 polynucleotide kinase (NEB). Radiolabeled transcripts were purified on polyacrylamide gel (6% acrylamide:bisacrylamide 19:1, 8 M urea).

EMSA were performed as previously described^60^. To determine binding affinity of Hfq to RyhB, ChiX and *ptsG*, radiolabeled RNA was heated for 1 min at 90 °C and put on ice for 1 min. RNA was diluted to 20 nM in modified Binding Buffer 2 (10 nM Tris-HCl pH 8.0, 1 mM DTT, 1 mM MgCl_2_, 20 mM KCl, 10 mM Na_2_HPO4-NaH_2_PO_4_ pH 8.0, 12.5 μg/mL yeast tRNA) and mixed with specific concentrations of Hfq (0-200 nM). Samples were incubated for 15 min at 37 °C and reactions were stopped by addition of 1 μL of non-denaturing loading buffer (1X TBE, 50% glycerol, 0.1% bromophenol blue, 0.1% xylene cyanol). For competition assays, 20 nM of radiolabeled *ptsG* was first incubated for 15 min at 37 °C with 100 nM Hfq (as described above). Specific concentrations of RyhB or ChiX (0-100 nM) were added to the samples and incubation was carried out for 15 min at 37 °C. Reactions were stopped by addition of 1 μL non-denaturing loading buffer. Sample were loaded on native polyacrylamide gels (5% acrylamide:bisacrylamide 29:1) in cold TBE 1X and migrated at 50 V, at 4 °C. Gels were dried and exposed to phosphor storage screens and analyzed using a Typhoon Trio (GE Healthcare) instrument. When applicable, quantification was performed using the Image studio lite software (LI-COR) and data was fitted using non-linear regression (GraphPad Prism).

### Fluorescence in situ hybridization (FISH)

Sample preparation for fixed cells was performed mostly according to the protocol previously reported^61, 62^. Briefly, ∼10 mL of cells was collected and fixed with 4% formaldehyde in 1x PBS for 30 minutes at room temperature (RT). The fixed cells were then permeabilized with 70% ethanol for 1 hour at RT. Permeabilized cells can be stored in 70% ethanol at 4 °C until the sample preparation. FISH probes were designed and dye-labeled as in the previous report^61, 62^. Hybridization was performed in 20 μL of hybridization buffer (10% dextran sulfate (Sigma D8906) and 10% formamide in 2x SSC) containing specific sets of FISH probes at 30 °C in the dark overnight. The concentration of FISH probes was 50 nM. After hybridization, cells were washed three times with 10% FISH wash buffer (10% formamide in 2x SSC) at 30 °C.

### Live-cell single-particle tracking and fixed cell SMLM imaging

Imaging was performed on a custom built microscopy setup previously described^63^. Briefly, an inverted optical microscope (Nikon Ti-E with 100x NA 1.49 CFI HP TIRF oil immersion objective) was fiber-coupled with a 647 nm laser (Cobolt 06-01), a 561 nm laser (Coherent Obis LS) and a 405 nm laser (Crystalaser). A common dichroic mirror (Chroma zt405/488/561/647/752r-UF3) was used for all lasers, but different emission filters were used for different fluorophores (Chroma ET700/75M for Alexa Fluor 647 and Chroma ET595/50M for mMaple3). For imaging Hoechst dye, a LED lamp (X-Cite 120LED) was coupled with a filter cube (Chroma 49000). The emission signal was captured by an EMCCD camera (Andor iXon Ultra 888) with slits (Cairn OptoSplit III), enabling fast frame rates by cropping the imaging region. During imaging acquisition, the Z-drift was prevented in real time by a built-in focus lock system (Nikon Perfect Focus).

For live-cell single particle tracking, 1 mL of cell culture was centrifuged by 1,500*g* for 5 min and 970 µL of the supernatant was removed. The remaining volume was mixed well and ∼1.5 µL was covered by a thin piece of 1% agarose gel on an ethanol-cleaned-and-flamed coverslip sealed to a custom 3D printed chamber. The agarose gel contained the same concentration of any drug or inducer used in each condition. Exceptions include rifampicin, which was at 100 μg/mL in the gel, due to high imaging background caused by high concentration of rifampicin, and IPTG for SP191 culture, which was eliminated in the gel, due to the high abundance mMaple3 already induced by IPTG in the culture. The power density of the 561 nm laser for single-particle tracking was ∼2750 W/cm^2^, and the power density of the 405 nm laser was ∼7 W/cm^2^ (except for SP191 where ∼4.5 W/cm^2^ was used due to high abundance of mMaple3). 1.5x Tube lens was used for the microscope body, and 2×2 binning mode was used for the camera. In this way, the effective pixel size became larger (173 nm instead of original 130 nm), receiving 77% more photons per pixel. 10 frames with 561 nm excitation were taken after each frame of 405 nm photo-conversion. About 13000 frames were collected per movie at a rate of 180 frames per second. For fixed-cell control experiment for tracking parameter optimization, imaging was performed using the exactly same imaging parameters as in live-cell measurements for a fair comparison. In cases imaging DNA together, Hoechst dye (Thermo 62249) was added to the ∼30 µL of cell culture before imaging as ∼20 µM final concentration, and imaged by the LED lamp (12%) with 500 ms exposure time. Imaging acquisition was conducted by NIS-Element (Nikon) software, at RT.

### Image reconstruction

The SMLM images are reconstructed as previously descried^62^, by a custom code written in IDL (Interactive Data Language). Briefly, all the pixels with an intensity value above the threshold were identified in each frame. The threshold was set at 3 times of the standard deviation of the individual frame pixel intensity. Among those pixels, the ones having larger values than surrounding pixels in each 5×5 pixel region are identified again as possible peak candidates, and 2D Gaussian function was fitted to a 15×15 pixel region surrounding these candidates. Candidates with failed fitting were discarded, and precise peak positions are defined for the remaining ones. The horizontal drift, which often occurs during the imaging acquisition, was corrected by fast Fourier transformation analysis.

### Tracking analysis

We used a MATLAB coded tracking algorithm to generate diffusion trajectories, which was modified by Sadoon and Yong^64^ based on the previously developed code^65^. Per each time step of ∼5.6 ms, 250 nm was empirically chosen to be the maximum one-step displacement to reduce artificial diffusion trajectories connected between different molecules, using a fixed cell sample as a control (Figure S3B). Trajectories longer than 5 time steps were used to calculate effective diffusion coefficient (D). Mean square displacement (MSD) as a function of time lag (Δt) was fit with a power law function (MSD = D(Δt)^α^). D values are reported in related figures. For one step displacement (*osd*) fitting, trajectories longer than 3 time steps were used, to populate many single *osd* values for fitting.

### Enrichment calculation

Enrichment at a certain region (nucleoid or cytoplasm) of a cell is defined as following.

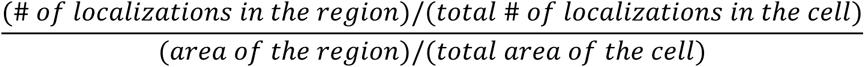

Here the ‘area’ of a cell refers to the two-dimensional area of the cell from the differential interference contrast (DIC) image (for the total area) or from the Hoechst image (for the nucleoid area), detected and calculated by our custom Matlab code^66^. “Cytoplasm” area/region is defined as the total region minus the nucleoid.

## Acknowledgements

We thank C.K. Vanderpool and X. Ma for sharing the plasmid containing mMaple3 template, E. Perozo for sharing the plasmid containing Lac repressor and operator system, Y. Wang for sharing MATLAB codes for extracting tracking trajectories, and the MRSEC Shared User Facilities at the University of Chicago (NSF DMR-1420709). We thank the Service de Purification de Protéines de l’Université de Sherbrooke (SPP) for Hfq purification and the Plateforme RNomique de l’Université de Sherbrooke for ddPCR. J. Fei acknowledges the support by the Searle Scholars Program, and NIH Director’s New Innovator Award (1DP2GM128185-01). Work in E. Massé Lab has been supported by an operating grant MOP69005 from the Canadian Institutes of Health Research (CIHR) and NIH Team Grant R01 GM092830-06A1.

## Author Contributions

J.F. conceived the project. J.F. and E.M designed the experiments. S.P., K.P. E.M.H. and M.C.C conducted the experiments, S.P., M.A.R., W.L. and J.F. analyzed the data, S.P. and J.F. interpreted the data, J.F. and E.M. wrote the manuscript.

