## Supplementary information for "Dynamic interactions between the RNA chaperone Hfq, small regulatory RNAs and mRNAs in live bacterial cells"

### Equal contribution

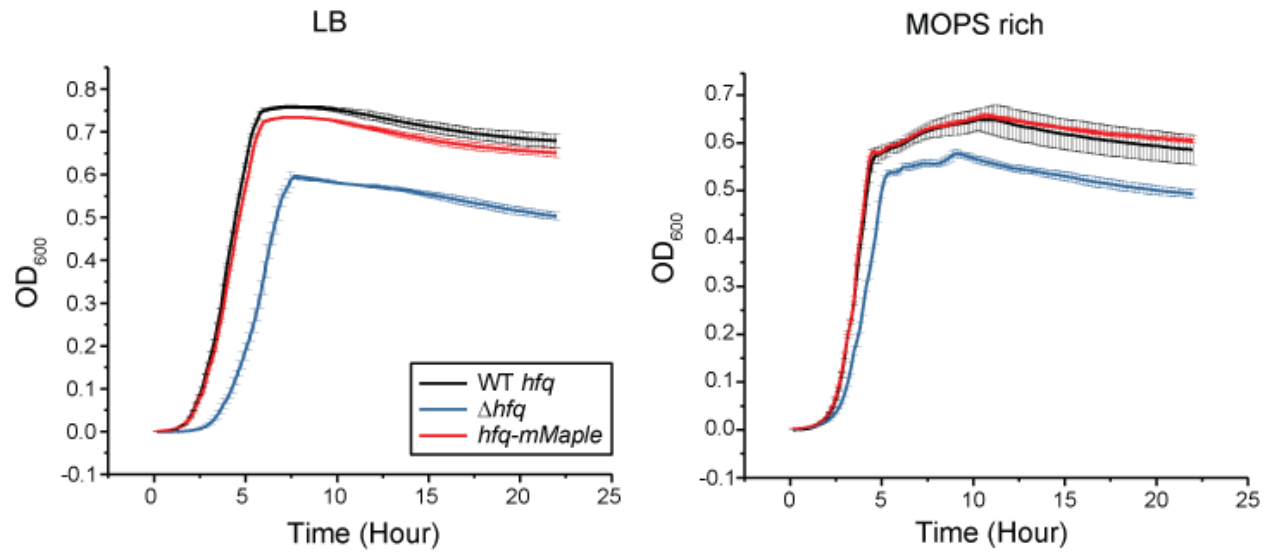

**Figure S1. mMaple3 tag on Hfq does not affect growth rate.** Growth curves of WT *hfq*, $\Delta hfq$ and *hfq-mMaple3* strains grown in LB or MOPS EZ-Rich containing 0.2% glucose medium. Data were obtained using a microplate spectrophotometer reader (BioTek). Error bars report the mean and standard deviation (s.d.) from 3 independent measurements.

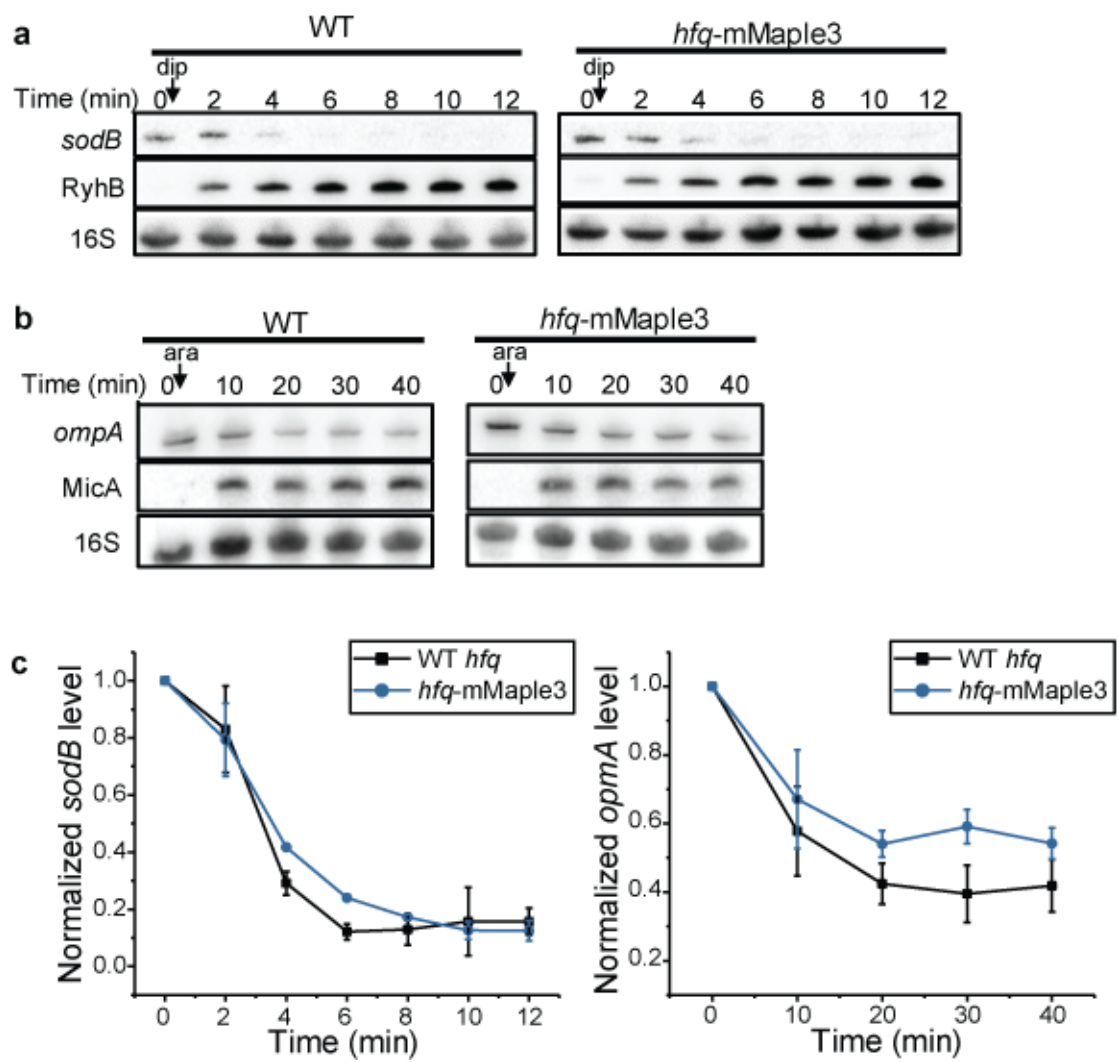

**Figure S2. mMaple3 tag on Hfq does not affect mRNA degradation by sRNA.** **a**, RyhB was induced by addition of 250 μM 2,2'-dipyridyl (dip) in a WT or in an *hfq-mMaple3* background when OD<sub>600</sub> reached 0.5 in LB medium. At indicated time points, total RNA was extracted. Specific *sodB* and RyhB probes were used for Northern blot analysis. **b**, MicA was induced by addition of 0.1% arabinose (ara) in a Δ*micA* or in a Δ*micA hfq-mMaple3* background, when OD<sub>600</sub> reached 0.5 in LB medium. At indicated time points, total RNA was extracted. Specific *ompA* and MicA probes were used for Northern blot analysis. For a and b, 16S rRNA was used as loading control. **c**, Densitometry analysis of *sodB* and *ompA* RNA levels obtained by Northern blots. Data was normalized to 16S at each time point to eliminate sample loading variation and normalized to the level at time 0 (before induction). Error bars report the mean and s.d. from 2-4 replicates.

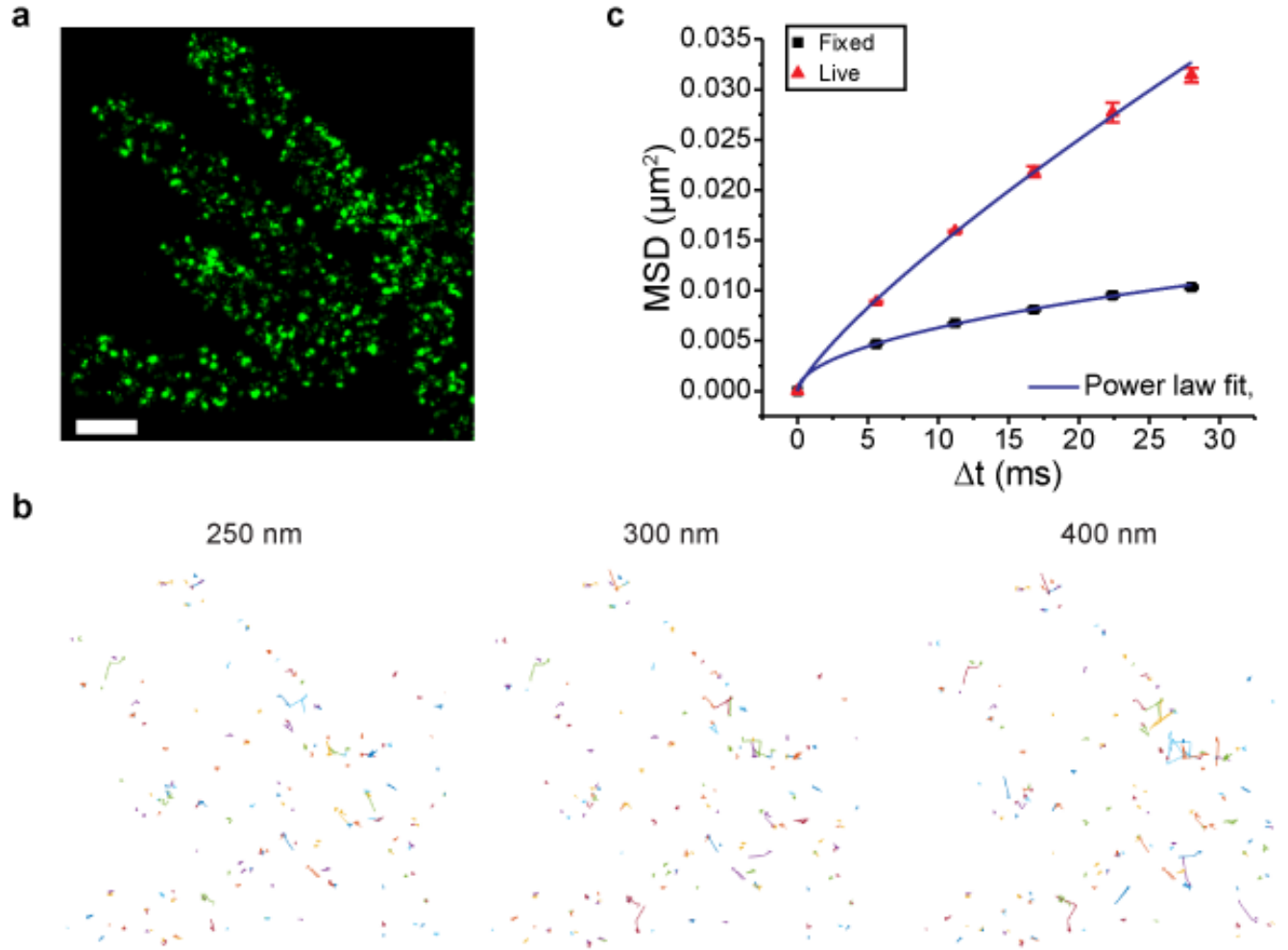

**Figure S3. Fixed cells as the stationary control for tracking analysis** **a**, A representative area with multiple fixed cells (*hfq-mMaple3* in WT *rne* background) under NT case is shown. **b**, Three examples of tracking by varying the distant threshold between neighboring frames. 250 nm gives minimal trajectories out of tiny clusters of spots, while 300 nm or 400 nm gives more and more artificial trajectories connecting different clusters. **c**, Mean square displacement (MSD) is plotted against time for the fixed cells, and power law fitting curves (blue) are shown.

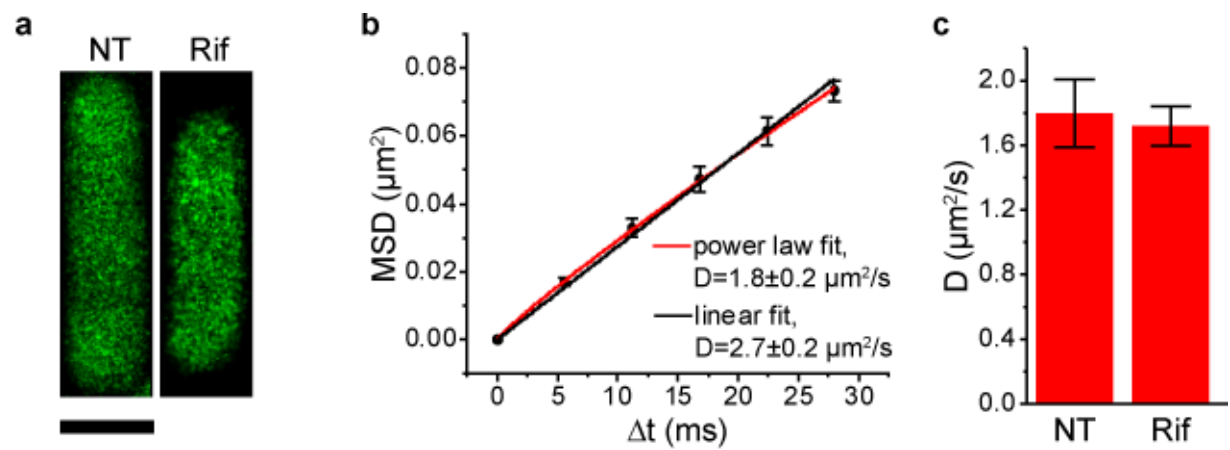

**Figure S4. Diffusivity of mMaple3 protein is not affected by treatment with rifampicin.** **a**, Representative images of control cells that express free mMaple3, under NT and Rif conditions. The scale bar represents 1 $\mu\text{m}$. **b**, Fitting with linear or power functions are demonstrated for one exemplary image area of mMaple3 under NT. **c**, Effective diffusion coefficients of mMaple3 from fitting with power function.

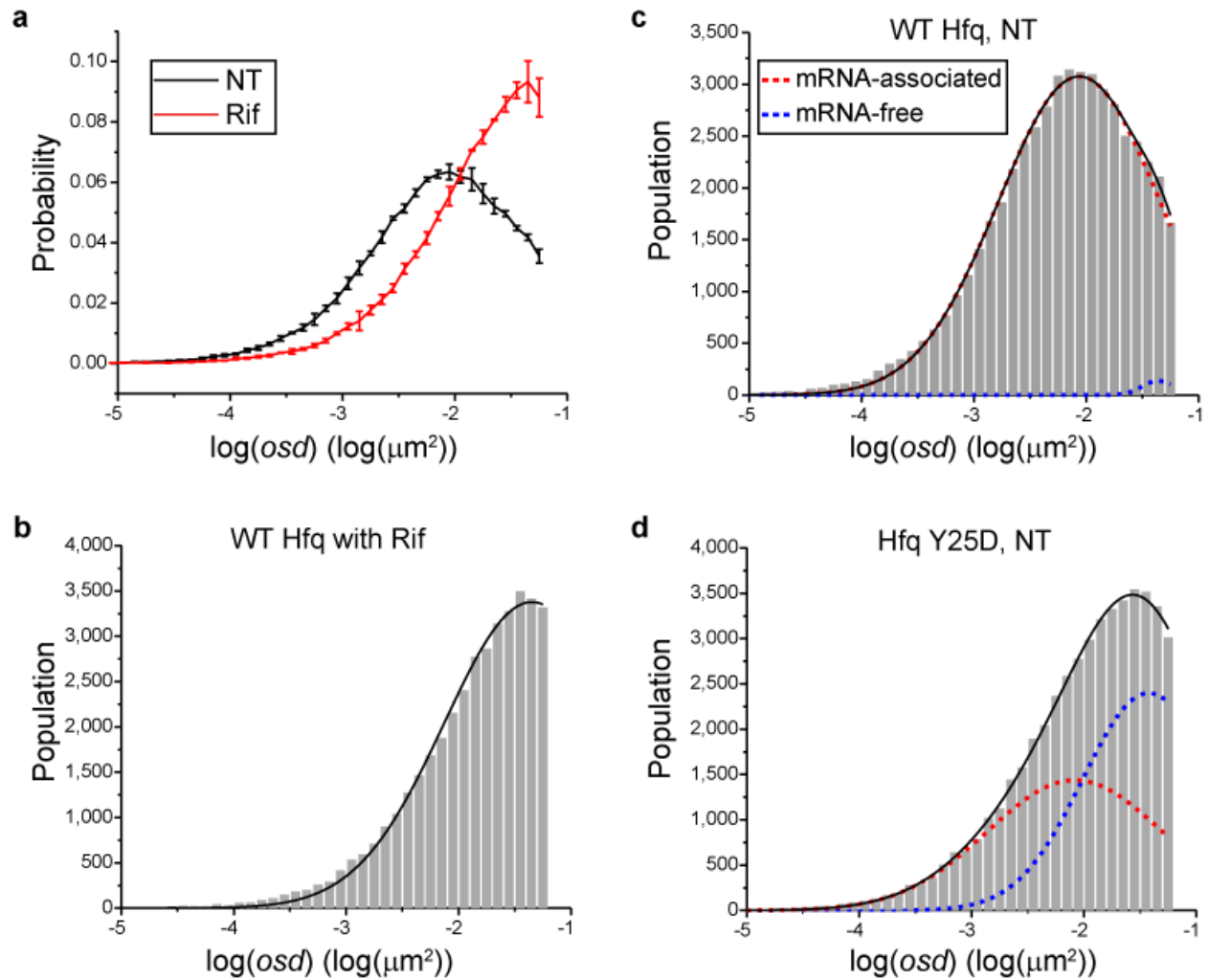

**Figure S5. Estimation of mRNA-associated Hfq fraction.** **a**, Normalized population distribution of *osd* for WT Hfq-mMaple3 under NT or rifampicin treatment. Error bars report s.d. from 2-3 replicates. **b**, An example of fitting *osd* distribution for WT under Rif with single Gaussian distribution (black curve). With the parameters constrained from b, Hfq-mMaple3 proteins under NT cases were fit with two Gaussian distributions. Blue and red curves represent the fitting of individual population, and black curves represent the overall fitting with two populations. The mRNA-associated fraction and mRNA-free fraction of Hfq proteins are estimated by the amplitudes of Gaussian peaks. Examples are shown for WT Hfq (**c**) and Hfq Y25D mutant (**d**).

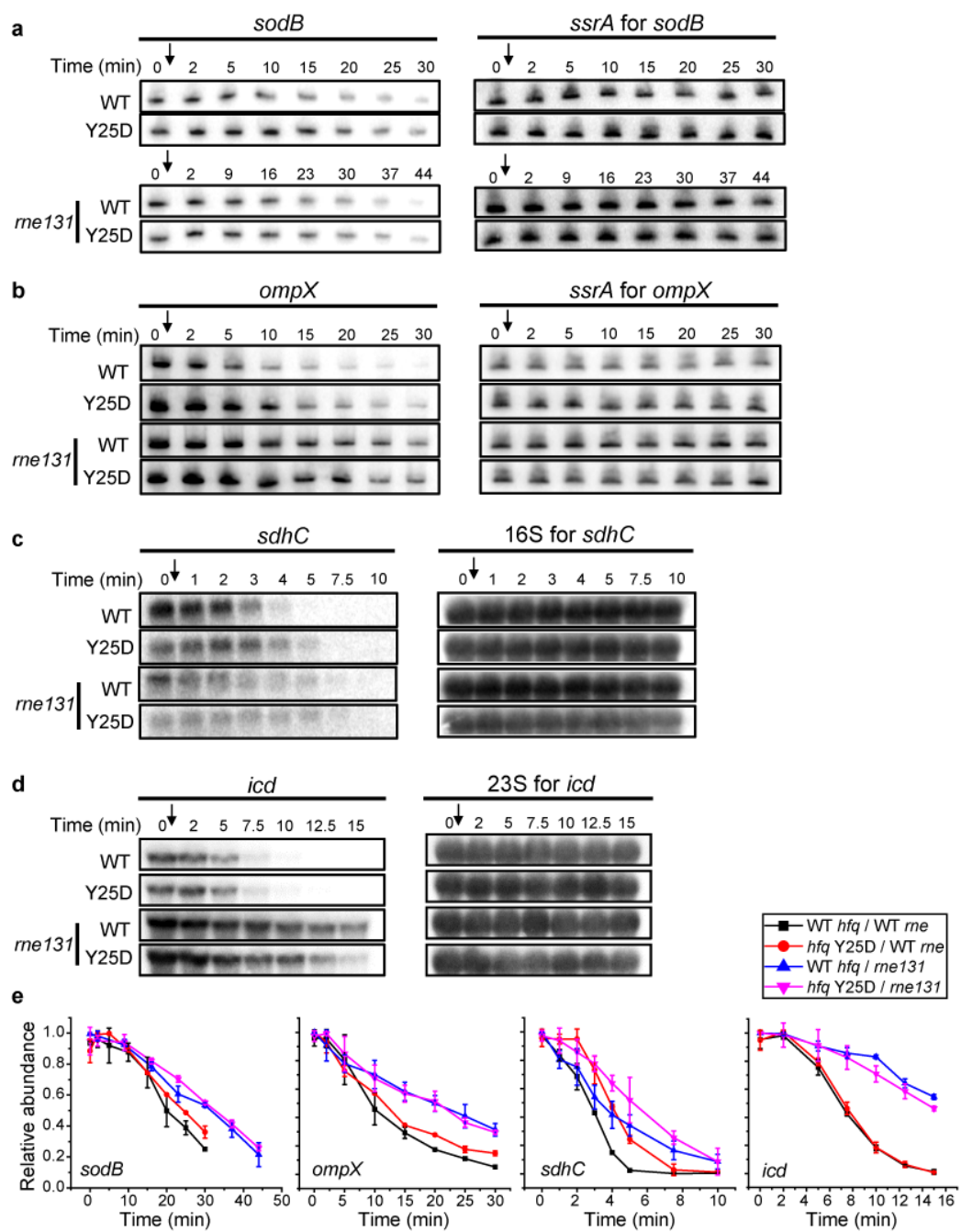

**Figure S6. mRNA half-life measurement by Northern blot.** **a**, Representative images of Northern blot. Strains were grown in MOPS EZ-Rich containing 0.2% glucose medium until $OD_{600} = 0.5$. Rifampicin (500 $\mu\text{g/mL}$) was added and total RNA was extracted at specific time points. Backgrounds: $\Delta\text{ryhB}\Delta\text{fnrS}$ (*sodB*), $\Delta\text{cyaR}\Delta\text{micA}$ (*ompX*), $\Delta\text{ryhB}\Delta\text{spf}\Delta\text{rybB}$ (*sdhC*). Specific *sodB*, *ompX*, *sdhC* and *icd* probes were used. **b,c,d**, *ssrA*, 16S rRNA or 23S rRNA were used as loading controls. **e**, Relative abundance of mRNA quantified by densitometry as a function of time. Each data point is represented by mean and s.d. of 2-3 biological replicates.

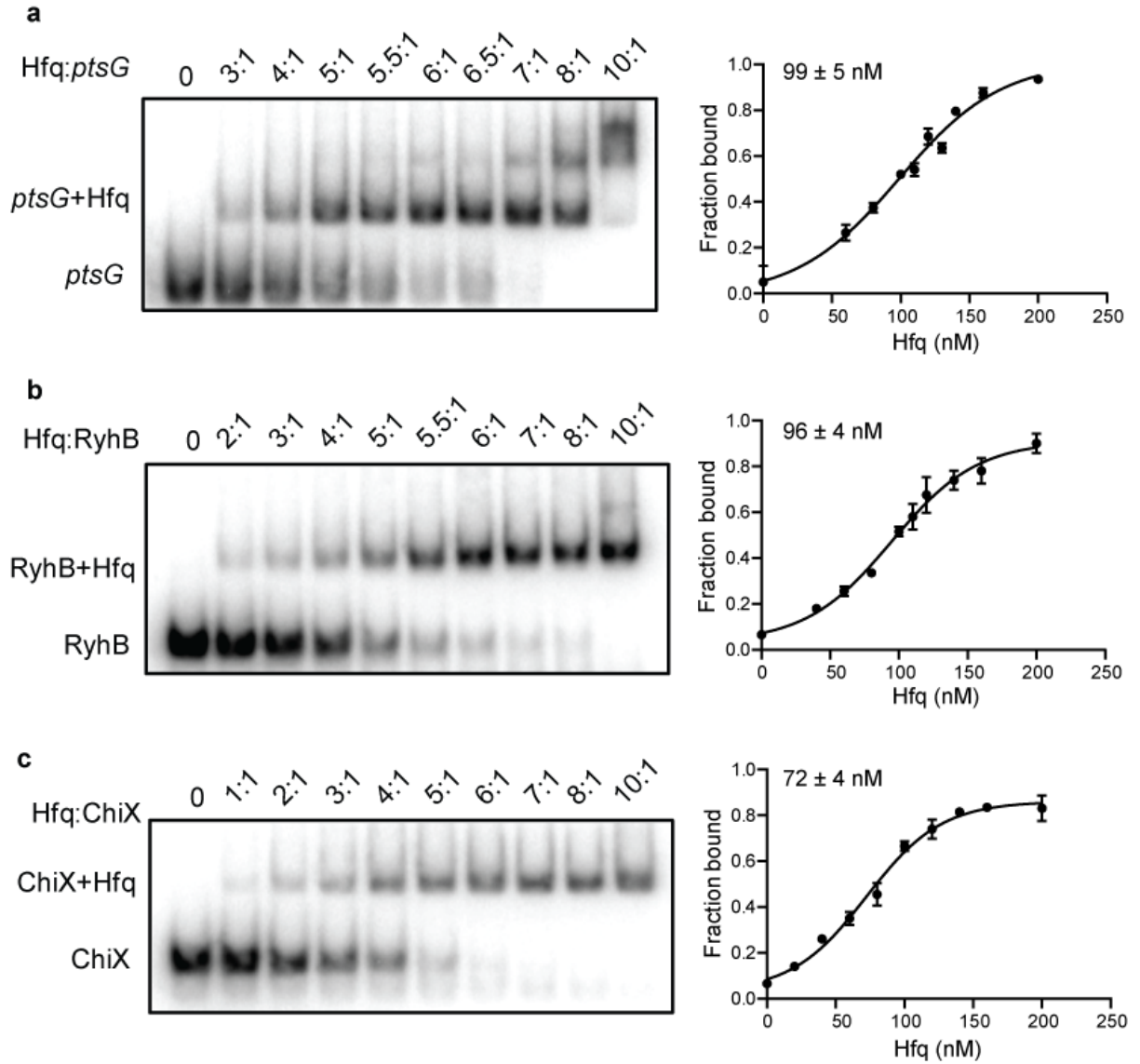

**Figure S7. Determination of $K_d$ values for Hfq interaction with RNAs.** 20 nM radiolabeled ($\gamma$) RNAs was incubated with increasing concentration of Hfq (20 - 2000 nM) **a**, γ-*ptsG*, **b**, γ-*RyhB*, **c**, γ-*ChiX*. Fraction bound was determined by densitometry of 2 independent experiments. To determined $K_d$, data was fitted using non-linear sigmoidal regression on GraphPad Prism.

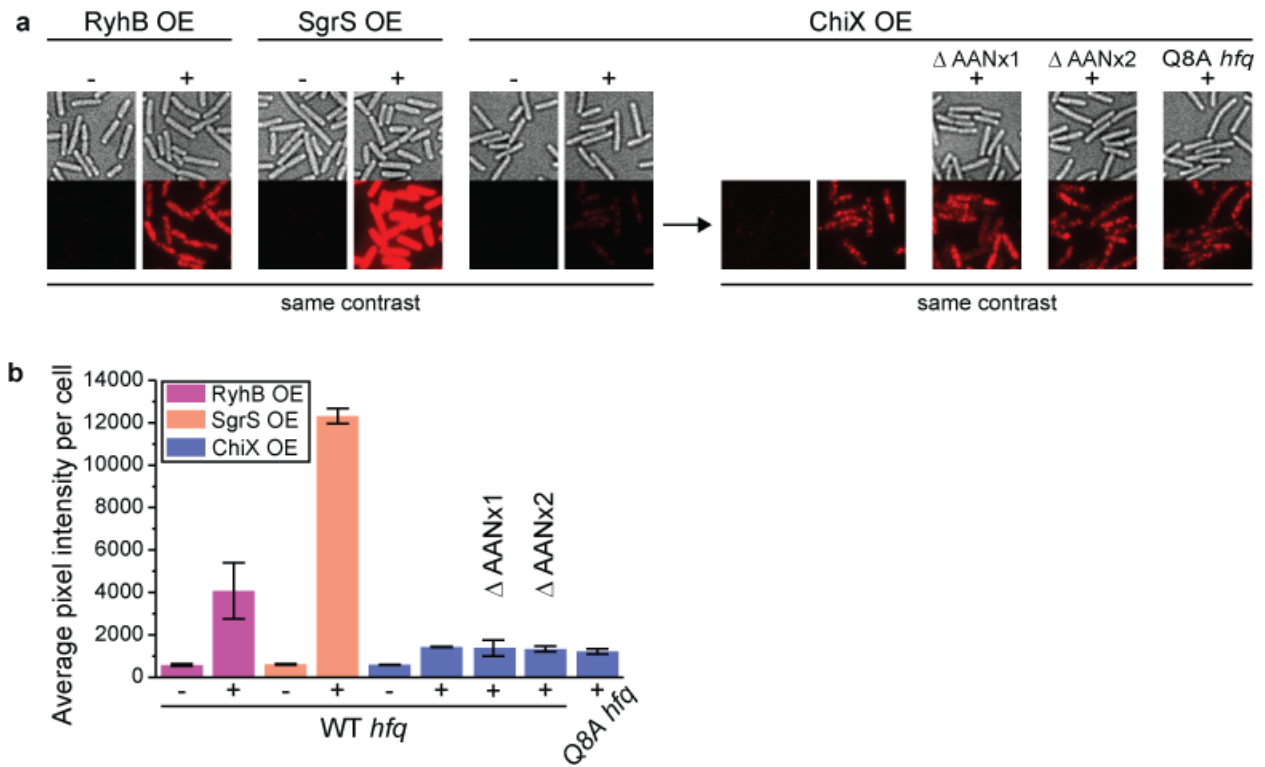

**Figure S8. FISH images and quantification of sRNA expression.** **a**, Representative FISH images of SgrS, RyhB, WT and mutant ChiX. Each sRNA was labeled with 3 FISH probes. Each sRNA was expressed from pET15b plasmid, induced by IPTG 1mM for 50 minutes (Table S1) in MOPS EZ defined medium. IPTG was added when OD<sub>600</sub> reached ~0.2. **b**, Quantification of expressed sRNAs. The staining efficiency of ChiX is the lowest, consistent with ChiX being more protected by Hfq.

**Figure S9. Uncropped northern blots and gel images.** The following images are uncropped northern blotting images used in main figures and supplementary figures. All blots were cropped at the indicated box.

**1. Uncropped gel images for EMSA of Figure 4c**

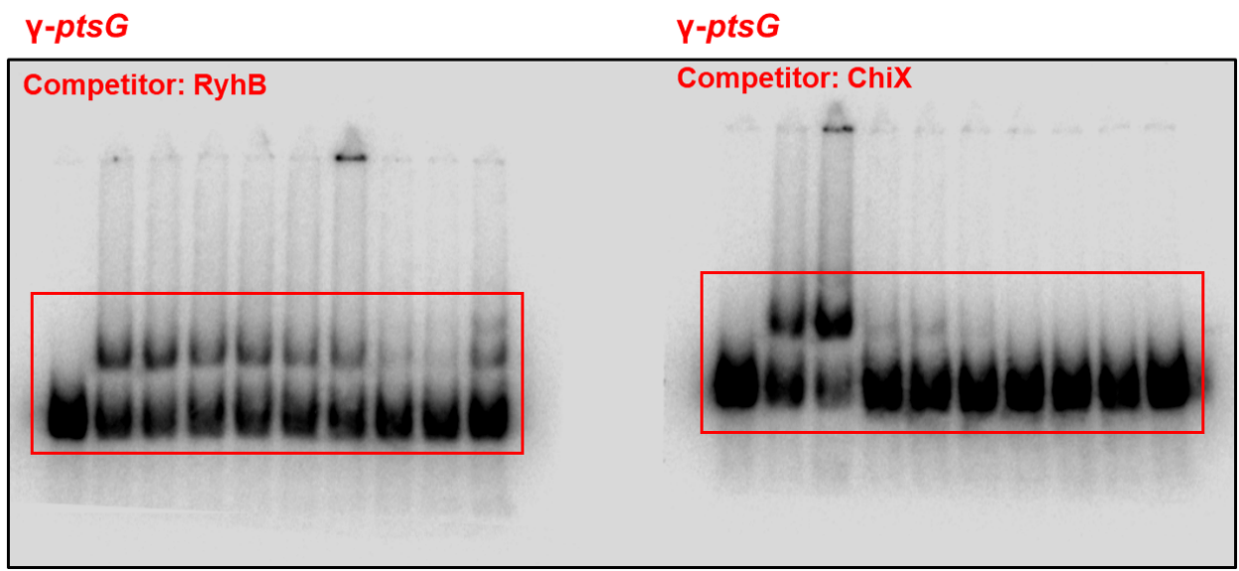

#### 2. <u>Uncropped Northern blot images of Figure S2</u>

**a**

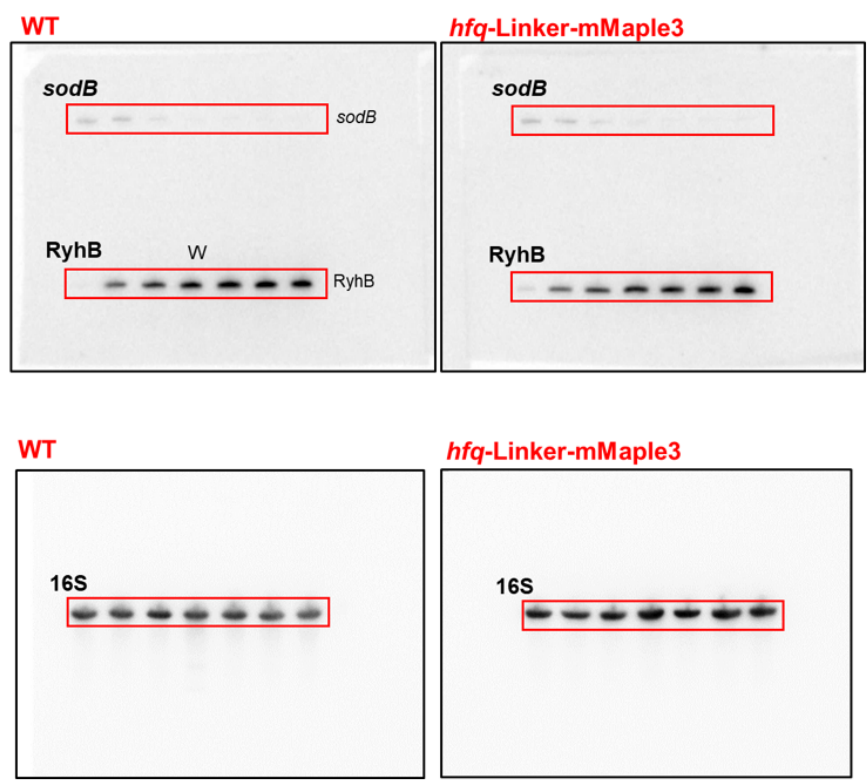

**b**

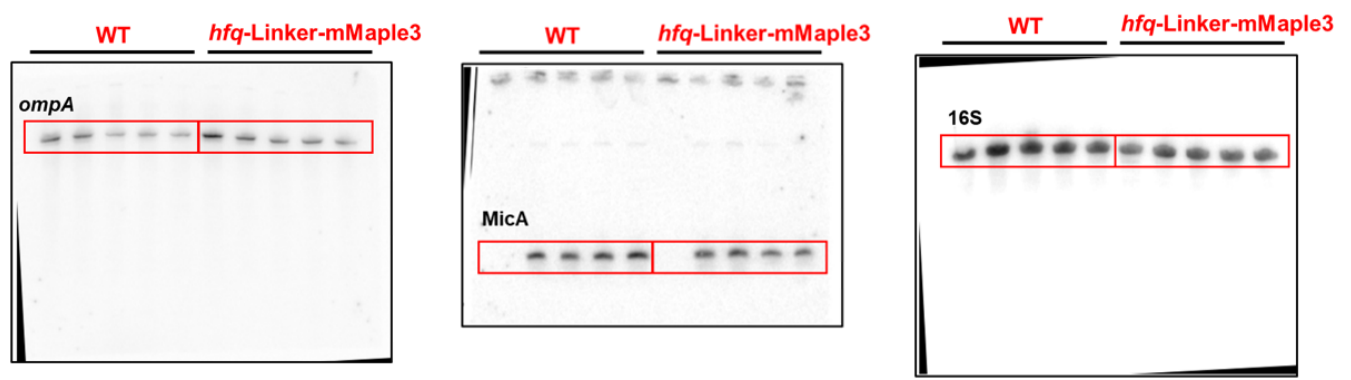

##### 3. <u>Uncropped Northern blot images of Figure S6</u>

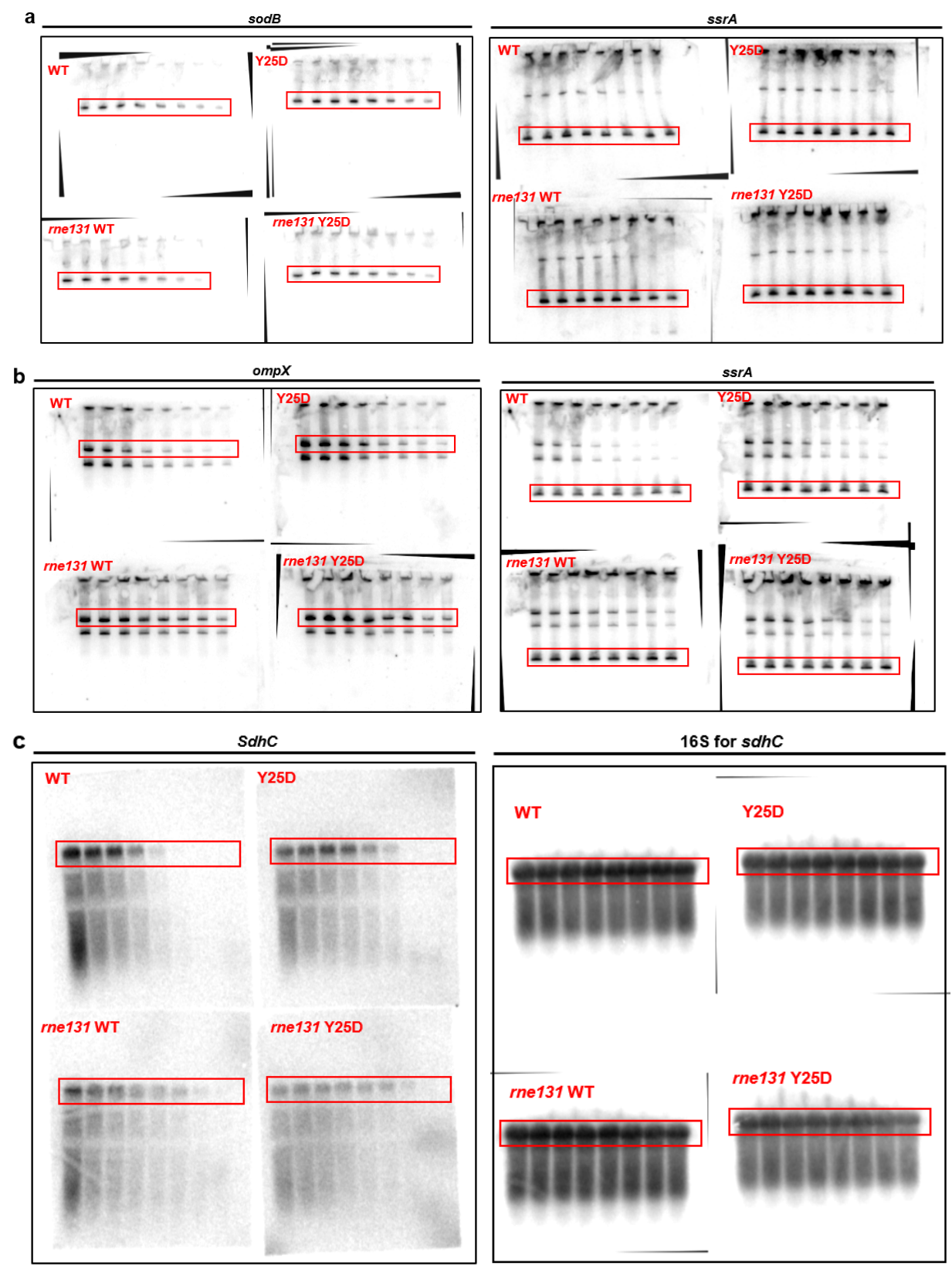

d

*icd*

23S

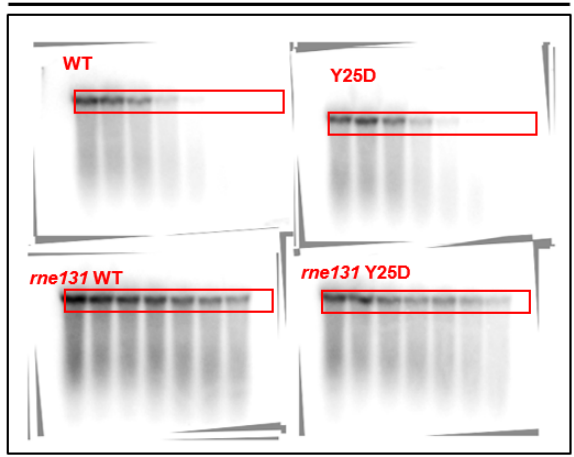

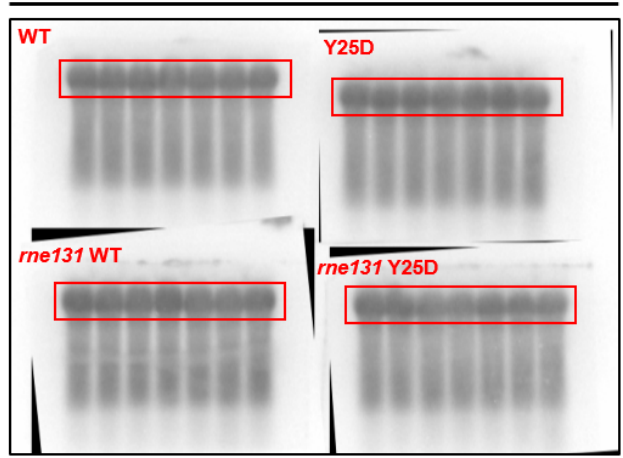

###### 4. Uncropped EMSA gel images for $K_d$ measurement for Figure S7

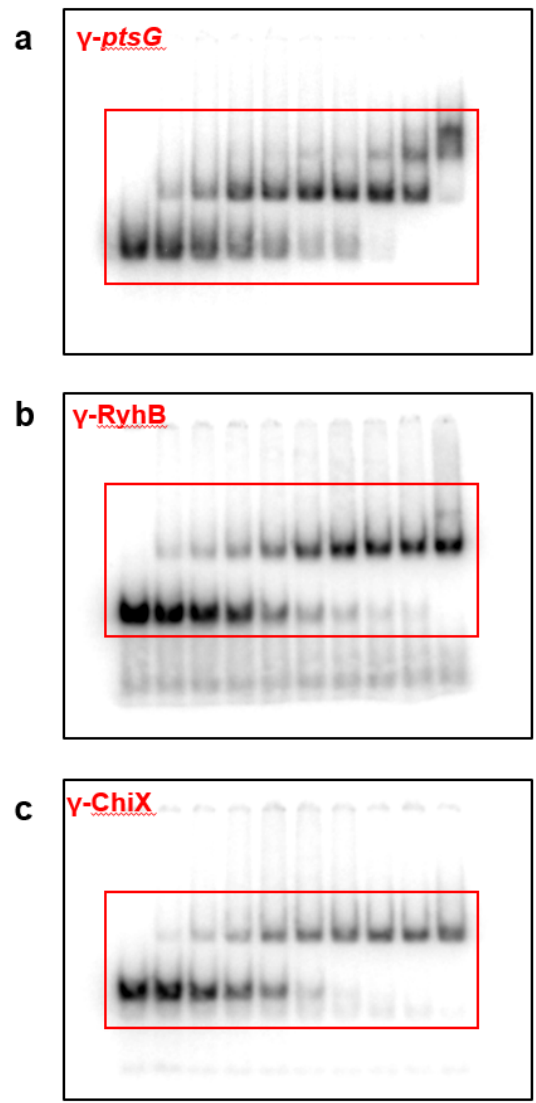

**Table S1.** List of all strains and plasmids used in this study

| Strains | Description | Reference |
| --- | --- | --- |
| AZZ41 | MC4100; *hfq*K31A | Gift from S. Gottesman Lab |
| DB166 | Δlac X74, *laq*, *tetR*, spec in MG1655 background | Thairu et al. 2017 |
| DL527 | EM1055 Δ*rybB* | Lalaouna et. al., 2015 |
| DL903 | EM1055 Δ*micA*::*cat* | This study (EM1055 + p1MPB57) |
| DL991 | EM1237; Δ*cyaR*::*kan* | Lalaouna et. al, 2018 |
| DL999 | EM1237 Δ*fnrS*::*kan* | This study |
| DL1029 | EM1055 Δ*cyaR*::*kan* | Lalaouna et. al, 2018 |
| DL1037 | EM1055 Δ*fnrS*::*kan* | This study (EM1055 + P1DL999) |
| DL1071 | Δ*ara714 leu+*; Δ*micA*::*cat* | This study (EM1451 + P1DL903) |
| EM1055 | MG1655 *lacX74* (WT) | Masse and Gottesman, 2002 |
| EM1237 | DY330 [W3110 delta-lacU169 gal490 lambda-cl857 delta-(cro-bioA)] | Yu et al., 2000 |
| EM1238 | EM1055 Δ*ryhB*::*cat* | Masse and Gottesman, 2002 |
| EM1376 | EM1055 *rne*Δ14 *zce*-726::Tn10 | Masse, Escorcia and Gottesman 2003 |
| EM1377 | EM1055 *rne*131 *zce*-726::Tn10 | Masse, Escorcia and Gottesman 2003 |
| EM1392 | BL21 DE3 pLysS; p15A CmR | Sigma |
| EM1451 | EM1055 Δ*ara714 leu+* | Desnoyers et. al., 2009 |
| GD549 | EM1055 Δ*rybB*::*kan* | Desnoyers and Massé, 2012 |
| JF133 | *rne*131 *zce*-726::Tn10; Δ*ryhB*::*cat* | Prévost et. al., 2011 |
| JW4130-1 | *rrnB3* ΔlacZ4787 *hsdR514* $\Delta$(*araBAD*)567 $\Delta$(*rhaBAD*)568 *rph*-1 Δ*hfq*-722::*kan* | Baba et al. 2006 |
| KK2560 | MG1655 *hfq*Q8A | Gift from S. Gottesman Lab (Zhang et al., 2013) |
| KK2561 | MG1655 *hfq*R16A | Gift from S. Gottesman Lab (Zhang et al., 2013) |
| KK2562 | MG1655 *hfq*Y25D | Gift from S. Gottesman Lab (Zhang et al., 2013) |
| KP1224A | EM1055; Δ*hfq*::*kan* | This study (EM1055 + P1JW4130-1) |
| KP1238 | EM1055 Δ*spf*::*kan* | This study (EM1055 + P1KP1237) |
| KP1867 | EM1055 *hfq*-Linker-mMaple3 linked to Kan | This study |
| KP1948A | EM1055 *hfq*Y25D-Linker-mMaple3 linked to Kan | This study |
| KP1952A | EM1055 *hfq*Q8A-Linker-mMaple3 linked to Kan | This study |
| KP1953 | EM1055 *hfq*R16A-Linker-mMaple3 linked to Kan | This study |
| KP1954 | *rne*131 *zce*-726::Tn10; *hfq*-Linker-mMaple3 linked to Kan | This study |
| KP1955 | *rne*131 *zce*-726::Tn10; *hfq*Y25D-Linker-mMaple3 linked to Kan | This study |
| KP1956 | *rne*131 *zce*-726::Tn10; *hfq*Q8A-Linker-mMaple3 linked to Kan | This study |
| KP1957 | *rne*131 *zce*-726::Tn10; *hfq*R16A-Linker-mMaple3 linked to Kan | This study |
| KP2044 | *rne*Δ14 *zce*-726::Tn10; *hfq*-Linker-mMaple3-Kan | This study (EM1376 + P1KP1867) |
| KP2045 | *rne*Δ14 *zce*-726::Tn10; *hfq*Y25D-Linker-mMaple3 linked to Kan | This study (EM1376 + P1KP1948A) |
| KP2046 | *rne*Δ14 *zce*-726::Tn10; *hfq*Q8A-Linker-mMaple3 linked to Kan | This study (EM1376 + P1KP1952A) |
| KP2047 | *rne*Δ14 *zce*-726::Tn10; *hfq*R16A-Linker-mMaple3 linked to Kan | This study (EM1376 + P1KP1953) |
| KP2136 | EM1055 Δ*cyaR* | This study |
| KP2137 | EM1055 Δ*ryhB*::*cat*; Δ*fnrS*::*kan* | This study (EM1238 + P1DL1037) |
| KP2140 | *rne*131 *zce*-726::Tn10; Δ*cyaR*::*kan* | This study (EM1377 + P1DL1029) |
| KP2141 | EM1055 $\Delta r y h B::cat$; $\Delta f n r S$ | This study |
| KP2142 | *rne131 zce-726::Tn10*; $\Delta r y b B$ | This study |
| KP2143 | *rne131 zce-726::Tn10*; $\Delta c y a R$ | This study |
| KP2145 | EM1055 $\Delta c y a R$; $\Delta m i c A::cat$ | This study (KP2136 + P1DL903) |
| KP2147 | EM1055 $\Delta r y h B::cat$; $\Delta f n r S$; *hfq*-Linker-mMaple3 linked to Kan | This study (KP2141 + P1KP1867) |
| KP2148 | EM1055 $\Delta r y h B::cat$; $\Delta f n r S$; *hfq*Y25D-Linker-mMaple3 linked to Kan | This study (KP2141 + P1KP1948) |
| KP2149 | $\Delta a r a 714 l e u+$; $\Delta m i c A::cat$; *hfq*-Linker-mMaple3 linked to Kan | This study (DL1071 + P1KP1867) |
| KP2155 | *rne131 zce-726::Tn10*; $\Delta c y a R$; $\Delta m i c A::cat$ | This study (KP2143 + P1DL903) |
| KP2158 | *rne131 zce-726::Tn10*; $\Delta r y b B$; $\Delta s p f::kan$ | This study (KP2142 + P1KP1238) |
| KP2159 | EM1055 $\Delta r y b B$; $\Delta s p f::kan$ | This study (DL527 + P1KP1238) |
| KP2162 | *rne131 zce-726::Tn10*; $\Delta c y a R$; $\Delta m i c A::cat$; *hfq*-Linker-mMaple3 linked to Kan | This study (KP2155 + P1KP1867) |
| KP2163 | *rne131 zce-726::Tn10*; $\Delta c y a R$; $\Delta m i c A::cat$; *hfq*Y25D-Linker-mMaple3 linked to Kan | This study (KP2155 + P1KP1948) |
| KP2167 | *rne131 zce-726::Tn10*; $\Delta r y b B$; $\Delta s p f$ | This study |
| KP2168 | EM1055 $\Delta r y b B$; $\Delta s p f$ | This study |
| KP2169 | EM1055; *hfq*R19D-Linker-mMaple3 linked to Kan | This study (EM1055 + KP2131B) |
| KP2172 | EM1055 $\Delta c y a R$; $\Delta m i c A::cat$; *hfq*-Linker-mMaple3 linked to Kan | This study (KP2145 + P1KP1867) |
| KP2173 | EM1055 $\Delta c y a R$; $\Delta m i c A::cat$; *hfq*Y25D-Linker-mMaple3 linked to Kan | This study (KP2145 + P1KP1948) |
| KP2174 | *rne131 zce-726::Tn10*; $\Delta r y b B$; $\Delta s p f$; $\Delta r y h B::cat$ | This study (KP2167 + P1EM1238) |
| KP2175 | EM1055 $\Delta r y b B$; $\Delta s p f$; $\Delta r y h B::cat$ | This study (KP2168 + P1EM1238) |
| KP2176 | BL21 DE3 pLysS; p15A CmR; $\Delta h f q::kan$ | This study (EM1392 + P1KP1224A) |
| KP2178 | *rne131 zce-726::Tn10*; $\Delta r y h B::cat$; $\Delta f n r S::kan$ | This study (JF133 + P1DL1037) |
| KP2179 | *rne131 zce-726::Tn10*; $\Delta r y h B::cat$; $\Delta f n r S$ | This study |
| KP2182 | EM1055 $\Delta r y b B$; $\Delta s p f$; $\Delta r y h B::cat$; *hfq*-Linker-mMaple3 linked to Kan | This study (KP2175 + P1KP1867) |
| KP2183 | EM1055 $\Delta r y b B$; $\Delta s p f$; $\Delta r y h B::cat$; *hfq*Y25D-Linker-mMaple3 linked to Kan | This study (KP2175 + P1KP1948) |
| KP2184 | *rne131 zce-726::Tn10*; $\Delta r y b B$; $\Delta s p f$; $\Delta r y h B::cat$; *hfq*-Linker-mMaple3 linked to Kan | This study (KP2174 + P1KP1867) |
| KP2185 | *rne131 zce-726::Tn10*; $\Delta r y b B$; $\Delta s p f$; $\Delta r y h B::cat$; *hfq*Y25D-Linker-mMaple3 linked to Kan | This study (KP2174 + P1KP1948) |
| KP2203 | EM1055 *hfq*K31A-Linker-mMaple3 linked to Kan | This study |
| KP2205 | EM1055; *hfq*K31A-Linker-mMaple3 linked to Kan | This study |
| KP2211 | *rne131 zce-726::Tn10*; $\Delta r y h B::cat$; $\Delta f n r S$; *hfq*-Linker-mMaple3 linked to Kan | This study (KP2179 + P1KP1867) |
| KP2212 | *rne131 zce-726::Tn10*; $\Delta r y h B::cat$; $\Delta f n r S$; *hfq*Y25D-Linker-mMaple3 linked to Kan | This study (KP2179 + P1KP1948) |
| KP2213 | EM1055 *hfq*F42A-Linker-mMaple3 linked to Kan | This study (EM1055 + P1KP2130A) |
| MPB57 | EM1237; $\Delta m i c A::cat$ | This study |
| SP191 | DB166 + pLac-mMaple3 | This study |

| Plasmids | Description | Reference |
| --- | --- | --- |
| pNM12 | pBAD24 derivative (arabinose inducible promoter; Amp<sup>R</sup>) | Majdalani et al., 1998 |
| pKD4 | Template for amplification of kanamycin resistance gene | Datsenko and Wanner, 2000 |
| pKD46 | Repts, Amp<sup>r</sup>, Rec recombinase expression vector | Datsenko and Wanner, 2000 |
| pZESA93M | Plasmid containing the *lac*operator and *mMaple3* gene | Gift from CK Vanderpool Lab |
| pLac-mMaple3 | Over expression of *mMaple3* through induction by IPTG | This study |
| pCP20 | Yeast Flp recombinase gene and FLP expression vector | Cherepanov and Wackernagel, 1995 |
| pBAD-*micA* | Over expression of *micA* through arabinose induction | This study |
| pBAD-*ryhB* | Over expression of *ryhB* through arabinose induction | Massé et.al., 2003 |
| pET21b-*hfq* | Over expression of *hfq* through induction by IPTG | Prévost et. al., 2007 |
| pET15b-RyhB | Over expression of RyhB through induction by IPTG | This study |
| pET15b-SgrS | Over expression of SgrS through induction by IPTG | This study |
| pET15b-ChiX | Over expression of ChiX through induction by IPTG | This study |
| pET15b-ChiX ΔAANx1 | Over expression of ChiX with one AAN site deleted, through induction by IPTG | This study |
| pET15b-ChiX ΔAANx2 | Over expression of ChiX with two AAN sites deleted, through induction by IPTG | This study |

**Table S2.** List of all oligonucleotides used in this study

| Primers | Sequence 5'-3' |  |
| --- | --- | --- |
| EM1690 | **ggatcgctggctccccgtgtaaaaaaacagcccgaaacctta**<u>catatgaatatcctccttag</u> | Rv 3'end *hfq* + P2 region (pKD4) |
| EM4293 | <u>gaagcagctccagcctacacctaattaagcttggctgcaggt</u> | Rv P1 region (pKD4) + mMaple3 |
| EM4313 | **gcagaatacttccgcgcaacaggacagcgaagaaaccgaat**<u>ctggtggcggaggctctg</u> | Fwd *hfq* + Linker |
| EM4314 | <u>tctggtggcggaggctctggtggcggaggcgtggttagcaagggcgaaga</u> | Fwd Linker + mMaple3 |
| EM88 | tgtaatacgactcactatagggcgatcaggaagaccctcgc | Fwd T7-*ryhB* |
| EM1978 | aaaaaaaaagccagcacccggctggc | Rv T7-*ryhB* |
| T7-ChiX(F) | taatacgactcactataggacaccgtcgcttaaagtgacgg | Fwd T7-*chiX* |
| T7-ChiX(R) | aaaaaaatggccaatatcgctattgg | Rv T7-*chiX* |
| T7-ptsG(F) | taatacgactcactataggcacagaattcataaataaagggcg | Fwd T7-*ptsG* |
| T7-ptsG(R) | aattgagagtgctcctgagtatgggt | Rv T7-*ptsG* |
| EM4704 | ctgcaagggcaaatcgagtcttttgatcaggccgtgatcctgttgaa | Fwd construct *hfq* F42A |
| EM4705 | <u>ctttacaagatccgttcctgaacgcactgcgtcgggaagatgttccag</u> | Fwd construct *hfq* R19D |
| EM2800 | gcgaagtcaataaaactcttaccattcagggcaatat**gtgtaggctggagctgcttc** | Fwd *fnrS* knock-out |
| EM2801 | ggactcttaaagggtagacgctgataaataacaggcaaa**catatgaatatcctccttag** | Rv *fnrS* knock-out |
| EM2596 | ctgaactctttcttcccaggcgagtctgagtatatgtgtaggctggagctgcttc | Fwd *micA* knock-out |
| EM2597 | gcggtgtggctggaaaaacacgcctgacagaaaagacatatgaatatcctccttag | Rv *micA* knock-out |
| EM2115 | gtgctttctgaactgaacaaaaaagagtaaagttagtcgcgtgtaggctggagctgcttc | Fwd *spf* knock-out |
| EM2075 | ggatcttttctttcgcccaataaaaaacgccccagtcattcatatgaatatcctccttag | Rv *spf* knock-out |
| EM123 | gtcacacg**catatg**gctaaggggcaatcttta | Fwd *hfq* - cloning in pET21b (**NdeI**) |
| EM124 | cgctcga**agc**ttttattcggtttcttcgctgtcctg | Rv *hfq* - cloning in pET21b (**HindIII**) |
| EM2651 | catag**catgc**aaaaaggccactcgtgagtggc | Fwd *micA* (**MscI**) |
| EM2652 | **ccat**gaaagacgcgcatttgttatcatcatc | Rv *micA* (**SphI**) |
| EH368 | <u>gatactcctctagcgatcaggaagaccctcgc</u> | pET15b + *ryhB* |
| EH369 | <u>ttcctgatcgctagaggagtatcttgttatccgctcacaatg</u> | pET15b + *ryhB* |
| EH270 | gtggataaattgagaacgaaagatcaaaaaaa**aaagccagcaccggct** | *ryhB* + endogenous terminator |
| EH271 | tttttgatctttcgttctcaatttatccac**ctgaaaggaggaactatatccggatatcc** | endogenous terminator + pET15b |
| EH305 | <u>atactcctcta</u>acaccgtcgcttaaagtgacgg | pET15b + *chiX* |
| EH306 | agcgacggtg**tagaggagtatctgttatccgctcacaatg** | pET15b + *chiX* |
| EH376 | <u>caaatgttgcgct</u>aaaaaaatggccaatatcgctattgg | *chiX* + endogenous terminator |
| EH375 | <u>cgqcaaatcccttctc</u>tgaaggaggaaactatatccggatatcc | pET15b + *chiX* endogenous terminator |
| EH362 | <u>gatactcctctagatgaagcaagggggtgcc</u> | <u>pET15b + sgrS</u> |
| EH363 | <u>ccttgcttcatctagaggagtatcttgttatccgctcacaatg</u> | <u>pET15b + sgrS</u> |
| EH374 | <u>ttttttattctcgccgcgctaaaaactgaaaggaggaactatatccggatatcc</u> | <u>pET15b + sgrS endogenous terminator</u> |
| EH373 | <u>gtttttagcgcggcgagaataaaaaaaaccagcaggtataatctgctgg</u> | <u>sgrS + endogenous terminator</u> |
| EH159 | aataaaaaaatgaaattcctctttgacg | ChiX ΔAANX1 |
| EH160 | atgccgtcactttaagcgac | ChiX ΔAAN |
| EH161 | aaaaaaatgaaattcctctttgacg | ChiX ΔAANX2 |

| FISH Probes | Sequence 5'- 3' |  |
| --- | --- | --- |
| RyhB_1 | gcgagggtcttcctgatcg | RyhB |
| RyhB_2 | atgtcgtgctttcaggttct | RyhB |
| RyhB_3 | ccagcacccggctggctaa | RyhB |
| SgrS_1 | cttaaccaacgcaaccagca | SgrS |
| SgrS_2 | catggtaatcggttgga | SgrS |
| SgrS_3 | gtcaactttcagaattgcgg | SgrS |
| ChiX_1 | cccgtcaaagaggaatttca | WT and mutant ChiX |
| ChiX_2 | cgtcactttaagcgacggtg | WT and mutant ChiX |
| ChiX_3 | aaatggccaatatcgctatt | WT and mutant ChiX |

| Northern Probes | Sequence 5'- 3' |  |
| --- | --- | --- |
| EM131 | <u>taatacgactcactatagggagattaccatacgaggactcctg</u> | Rv *sdhC* for RNA probe with <u>T7 promotor</u> |
| EM132 | tgttaatctggacctacagacc | Fwd *sdhC* for RNA probe |
| EM156 | <u>taatacgactcactatagggagacatatcggtcagttcagggtg</u> | Rv *icd* for RNA probe with T7 promotor |
| EM157 | aggcgagcgtaaaatctcctg | Fwd *icd* for RNA probe |
| EM1678 | gagcaatgtcgtgctttcaggttctccgcgagggtcttcctga | *ryhB* DNA probe |
| EM1692 | cagagcatctttagcatatggtagtgcaggtaattcgaatg | *sodB* DNA probe |
| EM2267 | ctgaaagtactttacaacccgaaggccttcttcatacacg | 16S DNA probe |
| EM2695 | ggctgttgtcttcttcatagcggtatttcaggttgaaacc | *ompX* DNA probe |
| EM3397 | gttttaacgcttcaacccca | *ssrA* DNA probe |
| EM2688 | gtgagtggccaaaatttcatctctgaattcagggatgatg | *micA* DNA probe |
| EM2703 | gcgaaaccagccagtgccactgcaatcgcgatagctgtc | *ompA* DNA probe |
| EM2747 | cccattcggaaatcgccggttataacggttcatatcacc | 23S DNA probe |

| Droplet Digital PCR | Sequence 5'- 3' |
| --- | --- |

|  |  |
| --- | --- |
| 16S_rRNA.ecoli.q.F2 | ggaataccggtggcgaaggc |
| 16S_rRNA.ecoli.q.R2 | ctccccaggcggtcgactta |
| chiX.ecoli.q.F2 | ggcgcttaaagtgacggcataa |
| chiX.ecoli.q.R2 | caatatcgctattggcccgtcaaa |
| EM2296 | gcgatcaggaagaccct |
| EM2297 | cccggctggctaagtaata |
| sgrS.qec.kp.F | agcgtcccacaacgattaac |
| sgrS.qec.kp.R | caccaatactcagtcacacatga |

|  |
| --- |
| Forward ddPCR 16S |
| Reverse ddPCR 16S |
| Forward ddPCR ChiX |
| Reverse ddPCR ChiX |
| Forward ddPCR RyhB |
| Reverse ddPCR RyhB |
| Forward ddPCR SgrS |
| Reverse ddPCR SgrS |

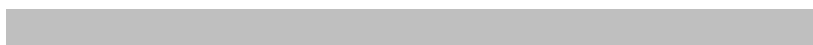

\_\_\_\_\_

\_\_\_\_\_

\_\_\_\_\_

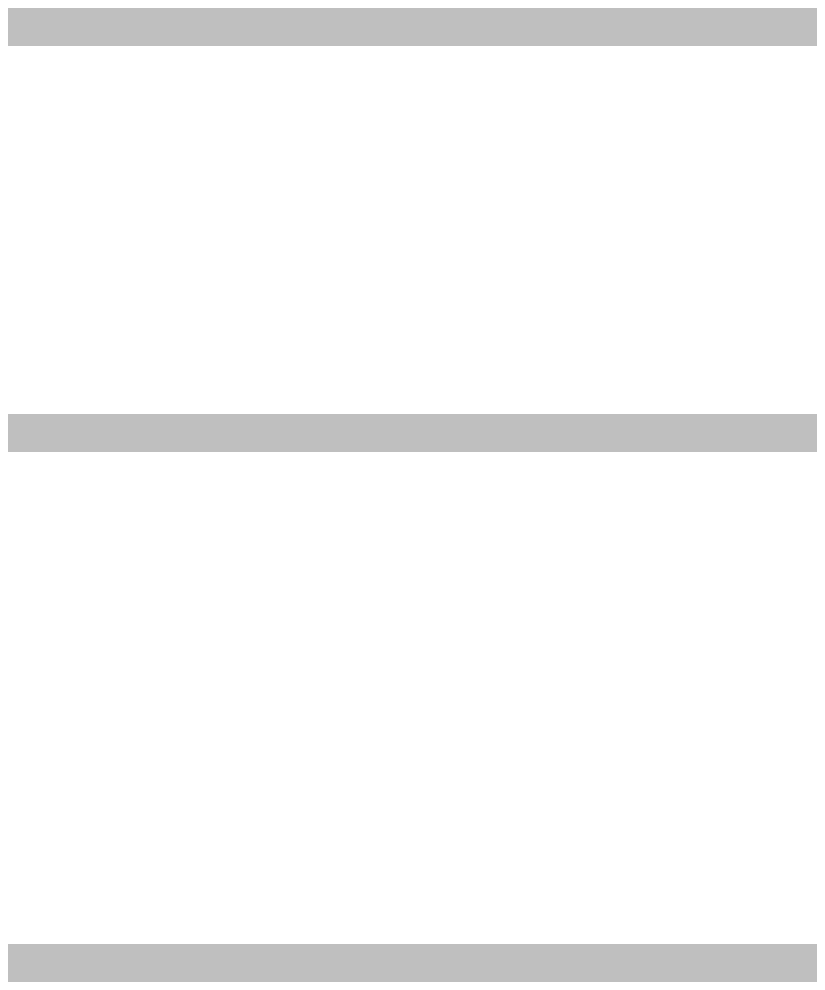
